# Conserved organ-specific microbial assemblages in different populations of a terrestrial crab

**DOI:** 10.1101/2021.03.30.437674

**Authors:** Giovanni Bacci, Sara Fratini, Niccolò Meriggi, Christine L. Y. Cheng, Ka Hei Ng, Massimo Pindo, Alessio Iannucci, Alessio Mengoni, Duccio Cavalieri, Stefano Cannicci

## Abstract

Brachyuran crabs originated in the oceans and evolved specific morphological and physiological adaptations to live in freshwater, intertidal and even terrestrial habitats but the role of a selection mechanism involving symbiotic microorganisms long these colonization processes are not known. In this work we investigated the associated microbiota of three populations of a terrestrial brachyuran crab, *Chiromantes haematocheir*, to find evidence of a conserved crab-specific microbiome unrelated to the population of origin and dissimilar from environmental microbial assemblages. Bacterial 16S rRNA gene and fungal ITS sequences were obtained from selected crab organs and environmental matrices to profile microbial communities. In spite of the presence of truly marine larval stages and no gregarious behaviour, favouring microbiota exchanges, we found common, organ-specific microbiota, associated to the gut and the gills of the crabs (with more than 15% of the genera detected specifically enriched only in one organ). Our results suggest an early establishment of a new common, stable microbiota in the transition from water to land.

## Introduction

The evolution of a terrestrial lifestyle independently occurred multiple times across and within most of the metazoan phyla *(1–4)*. Since the marine and terrestrial realms are ecologically very different, these several transitions involved profound morphological, physiological and behavioural adaptations, often converging towards similar structures and metabolic pathways *(1, 4)*. Among land adapted phyla, Arthropoda contribute the most to species diversity, which is the evolutionary outcome of a series of independent land invasions started in the Ordovician and still in progress at present *(4, 5)*. Indeed, insects and arachnids represent the most successful arthropods on land, in terms of biomass, functional roles and diversity, but crustaceans are also well represented. This largely marine group shows many examples of semi-terrestrial and terrestrial forms, mostly represented by Amphipoda, Isopoda and Decapoda, the latter including the largest extant terrestrial arthropod of the world, the iconic coconut crab *Birgus latro (6)*.

In chronological order, the true crabs (Decapoda; Brachyura) were among the last groups of arthropods to perform the sea-land transition *(5)*, but they have been particularly successful with more than twenty phylogenetically unrelated families that include species with intertidal, semiterrestrial and truly terrestrial lifestyles *(5, 7)*. Adaptations to terrestrial environments involved several morphological and physiological changes in sensory, locomotory, respiratory, excretory, osmoregulatory, digestive and reproductive systems *(8–18)*. Interestingly, brachyuran gills have not proved to be fully functional for respiration in terrestrial environment *(1)* and are involved in osmotic regulation, urine dilution, nitrogen excretion and ion-regulation in most terrestrially adapted families *(11, 19, 20)*. Other essential adaptations include herbivory and immune system recognition of potentially fungal pathogens, which are rare in marine environment but abundant on land, where they are commonly known pathogens of terrestrial arthropods, including semiterrestrial mangrove crabs *(21, 22)*.

Due to the relatively high diversity of terrestrial forms, when related to their recent history of attempts to conquer the land, brachyurans have been considered an excellent animal model for the study of the evolution of terrestrial adaptations *(2, 5, 8, 23)*. They are, however, rarely surveyed in studies investigating the involvement of commensal and symbiotic microbiota in such a dramatic transition. Indeed, metazoans groups have adapted to different habitats also by creating mutualistic and symbiotic relationships with the microorganisms that inhabit their tissues and organs *(24)*. These relationships can be so tight to drive the evolution and Darwinian selection of both host and microbiome, as a single unit *(25–27)*.

Within this theoretical frame, we investigated the role of associated microbial communities in the terrestrial adaptations of a semi-terrestrial brachyuran crab, *Chiromantes haematocheir*, a common inhabitant of lowland forests of East Asia *(28, 29)*. We applied targeted metagenomics approaches to profile both prokaryotic and fungal communities isolated from 1) organs (i.e., gills, gonads and gut) collected from specimens belonging to different populations and 2) relevant environmental matrixes (i.e., soil, leaf litter, water and water debris) sampled in their microhabitats. Our null hypothesis was that microbial composition of crab organs could show the same microbial signature displayed by the matrices from the surrounding environments. This would indicate that the crab organs would host bacterial and fungal populations acquired from the environments without any specific selection and could be explained by a dynamic “loss and acquire” balance that ultimately would prevent the development of stable host-microbe associations. Our alternative hypothesis was that crab organs may show exclusive bacterial and/or fungal communities, in terms of diversity and composition, when compared to those present in the environment. Thus, our goal was to investigate, for the first time for terrestrial brachyurans, the presence of a selection process in favour of specific microorganisms and, ultimately, the evolution of adaptive symbioses able to contribute to the terrestrial life style of a bimodal crab only recently adapted to the land environment.

## Results

### Microbial composition of crab’s organs and environmental samples

Sequencing produced 25,875,448 sequences for prokaryotic 16S rRNA amplicon and 15,788,842 sequences for the fungal internal transcribed region 1 (ITS1) that were clustered according to DADA2 pipeline (see Supplementary methods for additional details). After the clustering pipeline two fungal samples (namely GI_TKP_5 and MG_SH_1) produced no counts in any of the fungal amplicon sequence variants (ASVs) and were therefore removed from subsequent analyses. Since several fungal samples reported a low number of reads in crab’s organs, two samples were replicated and tested for correlation and accuracy to evaluate the presence of fungal cells in the crabs’ organs (Figure S1 and Table S1). Technical replicates showed an average correlation coefficient (Spearman’s rho) between replicates ranging between 0.74 and 0.99, with an accuracy higher than 75% in all contrasts. These results mean that the low recovery in biotic samples was not due to technical or analytical issues but corresponded to the scarce presence of fungal DNA (Figure S2, Table S2 and Supplementary method section).

Single nucleotide clustering detected a total of 56,233 ASVs including both 16S rRNA gene and ITS1 amplicons. The 16S rRNA gene amplicons produced 55,036 ASVs (97.87%), whereas the ITS1 region was clustered into 1,197 ASVs (2.13%). Most variants were assigned to the Bacteria domain (54,800 ASVs, 97.45% of the total ASVs detected) and Fungi kingdom, even if a sporadic presence of Archaea was found (236 ASVs, 0.42% of the total ASVS detected). In general, microbial diversity was higher in the environmental samples than in crab’s organs, except for the gills where we found a high diversity of 16S rRNA amplicons (Figure 2a and b, Table S3). The fungal diversity was low in all crab’s organs, whereas environmental samples were characterised by a highly diverse fungal assemblage, with soil, water, and water debris reporting the highest diversities (Figure 2c & d). For each sample category and at each site, the difference between extrapolated diversity—namely the inverse Simpson index calculated by simulating a higher sequencing depth than the observed depth for each sample—and observed diversity was low (Table 1), meaning that our sampling effort was effective in assessing the natural microbial diversity for each category. Even the Good’s coverage estimator was higher than 99.9% in all sample types highlighting that only 1‰ of clustered sequences came from ASVs detected only once in the whole dataset (Table 1 and Table S3). Microbial diversity was rather uniform across sampling sites, except for To Kwa Peng (TKP), which showed a significantly lower diversity with respect to the other sites in terms of ITS1 data (Figure S3).

**Table 1:** Mean differences between extrapolated diversity and observed diversity in each sample type and site. The differences between extrapolated Simpson diversity (Inverse Simpson index computed for a sequencing depth higher than the observed one) and observed diversity (Inverse Simpson index computed for a sequencing depth equal to the real sequencing depth of the sample) was reported using the average value ± the standard error on the mean for each sample type and site. Good’s coverage estimator was also reported using the same notation used for Simpson diversity. Abbreviations as in Figures 1 and 2.

|  |  | Inverse Simpson (ext - obs) | Good's coverage estimator |
| --- | --- | --- | --- |
| **Sample type** |  |  |  |
| | MG | 0.051 ± 0.040 | 99.995 ± 0.001 |
| | HG | 0.132 ± 0.126 | 99.997 ± 0.001 |
| | GO | 0.085 ± 0.029 | 99.987 ± 0.005 |
| | GI | 0.575 ± 0.230 | 99.994 ± 0.002 |
| | L | 0.185 ± 0.114 | 99.963 ± 0.007 |
| | S | 2.741 ± 0.809 | 99.993 ± 0.001 |
| | W | 0.246 ± 0.066 | 99.997 ± 0.001 |
| | WD | 1.149 ± 0.293 | 99.992 ± 0.002 |
| **Site** |  |  |  |
| | SH | 0.705 ± 0.236 | 99.987 ± 0.003 |
| | SK | 0.502 ± 0.146 | 99.990 ± 0.002 |
| | TKP | 0.240 ± 0.067 | 99.996 ± 0.001 |

**Figure 1.**
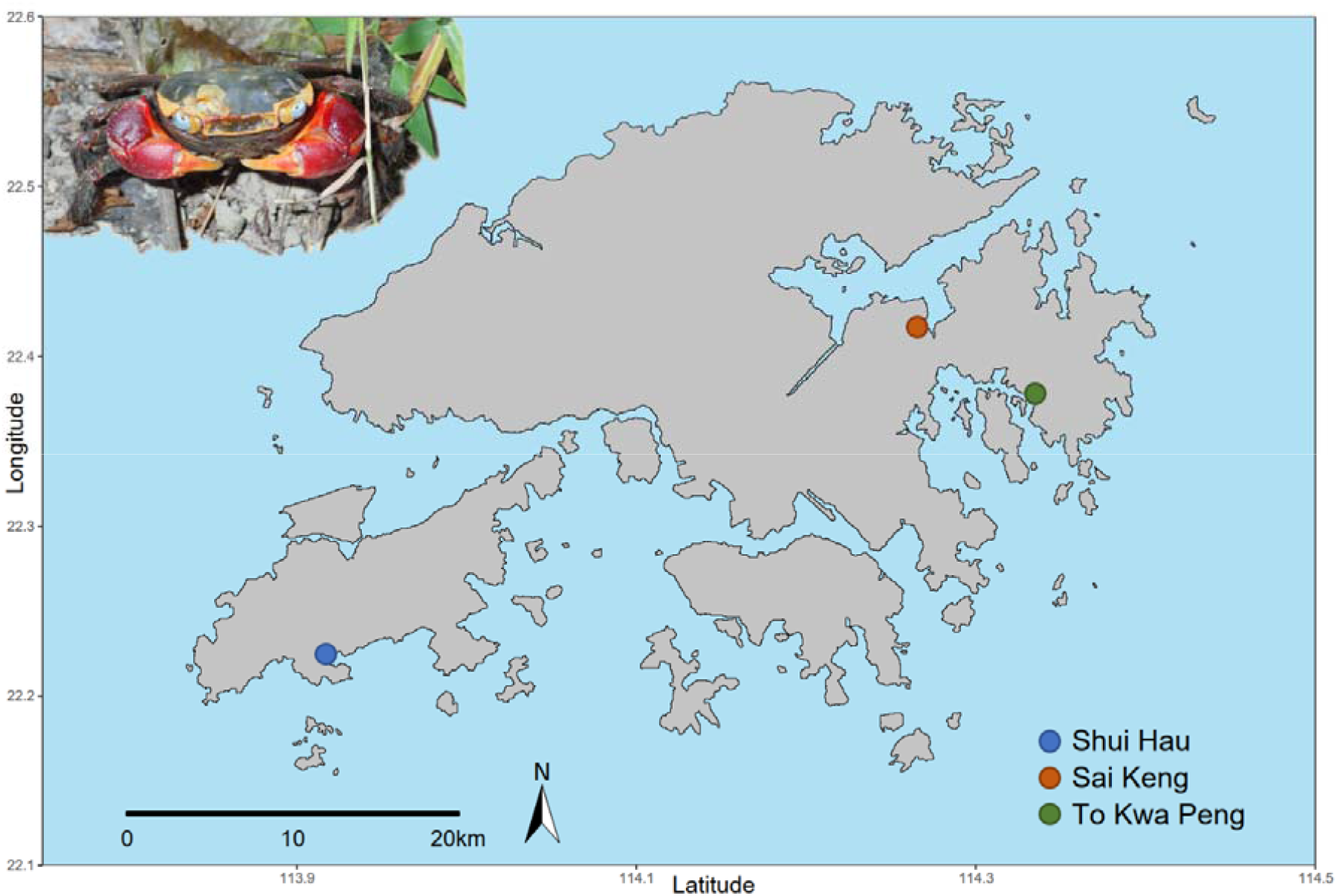
Geographical distribution of the sampled populations of *C. haematocheir*. The map shows the names and location of the three sampling sites visited for the study in the New Territories and on Lantau Island and an adult male *C. haematocheir* in its natural environment (top left).

**Figure 2:**
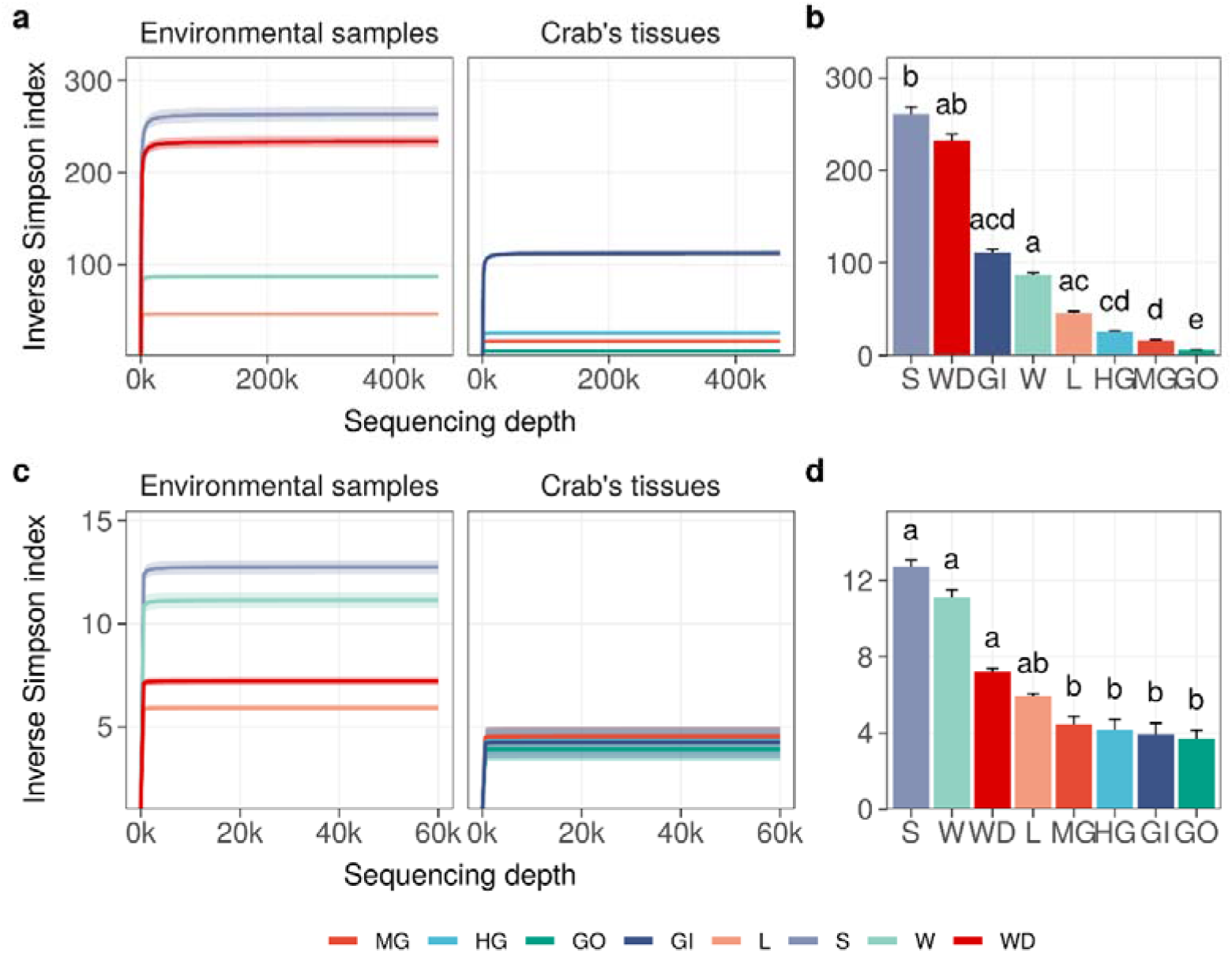
Microbial diversity in crab’s organs and environmental samples. The average of the inverse Simpson index was reported with increasing sampling effort for all types of samples. Interpolated and extrapolated diversity was reported in panel a and c (16S rRNA gene and ITS-1 region, respectively), whereas observed diversity was reported in panel b and d (16S rRNA gene and ITS-1 region respectively). Significant differences in microbial diversity (Wilcoxon non-parametric test) were reported using lowercase letters (panel b and d) whereas colours and acronyms on the x-axis correspond to different sample types (MG, midgut; HG, hindgut; GO, gonads; GI, gills; L, litter; S, soil; W, water; WD, water debris). If two means were significantly different, all letters on top of the two boxes must be different; if two means were equal, at least one letter must be the same.

### Microbial distribution across sample types

The multidimensional ordination showed that both environmental samples and crab’s organs contributed to shape microbial community distribution, but their effect varied according to both the amplicon type and the category of sample considered (Figure 3 and S4). In terms of composition, the microbial communities found in the environmental samples overlapped with each other more than the ones characteristic of the crab’s organs (Figure 3a). The PCoA built on 16S rRNA gene amplicons (Bacteria and Archaea) showed that soil and litter, and water and water debris, respectively, formed two distinct clusters. The observed intra-cluster variability is due to differences in microbial communities found across the three Hong Kong sites (Figure S4). The same 16S rRNA amplicon dataset showed a clear separation across crab’s organs, with a very limited influence of the collection sites (Figure 3 and S4). The gut’s microbiome composition was similar across the two sampled sections (mid- and hindgut), while the gills and gonads sharply separated from each other and the gut itself. These significant differences were highlighted also by the permutational analysis of variance (Figure 3b, Table S4). With respect to 16S rRNA data, fungal distribution produced more overlap, especially across crab’s organs, with the gonads being the only organ to show significant differences when compared to the other organs (Figure 3b, Table S4).

**Figure 3:**
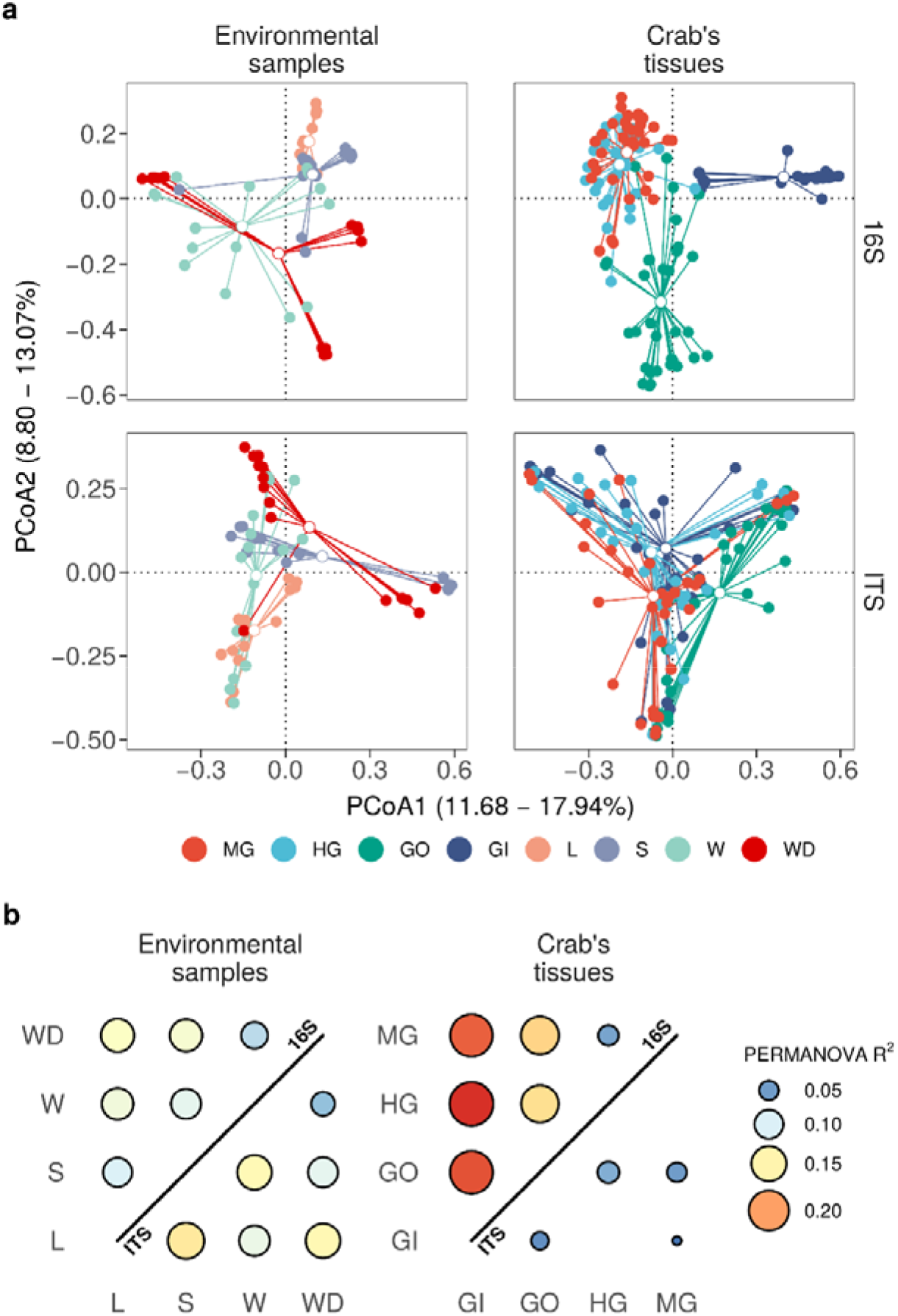
Microbial distribution according to Bray-Curtis distance. a) Principal coordinates analyses on Bray-Curtis distances inferred form 16S rRNA and ITS-1 metabarcoding. Environmental samples and crab’s organs (top side of the panels) were reported separately as distribution obtained with the two markers reported (right side of each panel). Solid-coloured points represent different samples whereas white-filled points represent centroids. The variance of the objects along each axis has been reported between squared brackets (from the lowest to the highest) and different sample types were reported using different colours (MG, midgut; HG, hindgut; GO, gonads; GI, gills; L, litter; S, soil; W, water; WD, water debris). b) Permutational analysis of variance on ordinations reported in panel a. For each pair of organs and environmental samples a permutational analysis of variance was performed. R-squared values of significant contrasts were reported for both 16S (upper triangle) and ITS (lower triangle) counts.

Both environmental samples and crab’s organs possessed a large set of microbial species which were unique to the category considered (Figure S5). As already mentioned, sampling sites also contributed to shape microbial communities, but they predominantly affected environmental samples distribution and showed a low effect on crab’s organs (Figure S4a and b, Figure S5a and c). In addition, the community composition of both prokaryotic and fungal communities was more variable and dispersed at site level than at organ level (Figures S4 and S6, Table S5), showing a higher specificity of such assemblages within the different organs and across sites. Gills exchanged microbes with environmental matrices in direct contact with them (soil and water) and even with the gut, which is in turn connected with the same environmental matrices through defecation (Figure S5b and d).

### Defining characteristic patterns across sample types

To inspect microbial distribution in different sample categories, we performed log-likelihood ratio test on both prokaryotic and fungal communities using DESeq2. We found 250 ASVs (218 bacteria and 32 fungi) reporting a different distribution across crab’s organs and/or environmental samples (Figure 4 and Table S6). Even if significant ASVs corresponded to 0.45% of the ASVs profiled in the whole community (250 on 55819), they accounted for more than 50% of the total microbial abundance (with a mean in each sample of 56.5% and a standard error of 2.10%) reflecting the presence of many rare and sporadic species throughout sample types and sites. Divisive analysis of hierarchical clustering obtained using variance-stabilized counts produced four distinct clusters, which show a peculiar pattern of abundance of ASVs clearly linked to the ecological and biological settings of the study. Indeed, clusters 1 and 2 (Figures 4 and S7) were composed by microbial assemblages highly represented in the crab’s organs, while the other two clusters, (namely, cluster 3 and 4) represented the microorganisms mostly found in the environmental samples (Figures 4 and S7). The clusters representing the microorganisms more abundant in the crab’s organs were represented by bacterial ASVs only and were split into two groups, formed by the bacteria of the gut (cluster 1) and of the gills (cluster 2), respectively, with the latter being the smaller one.

**Figure 4:**
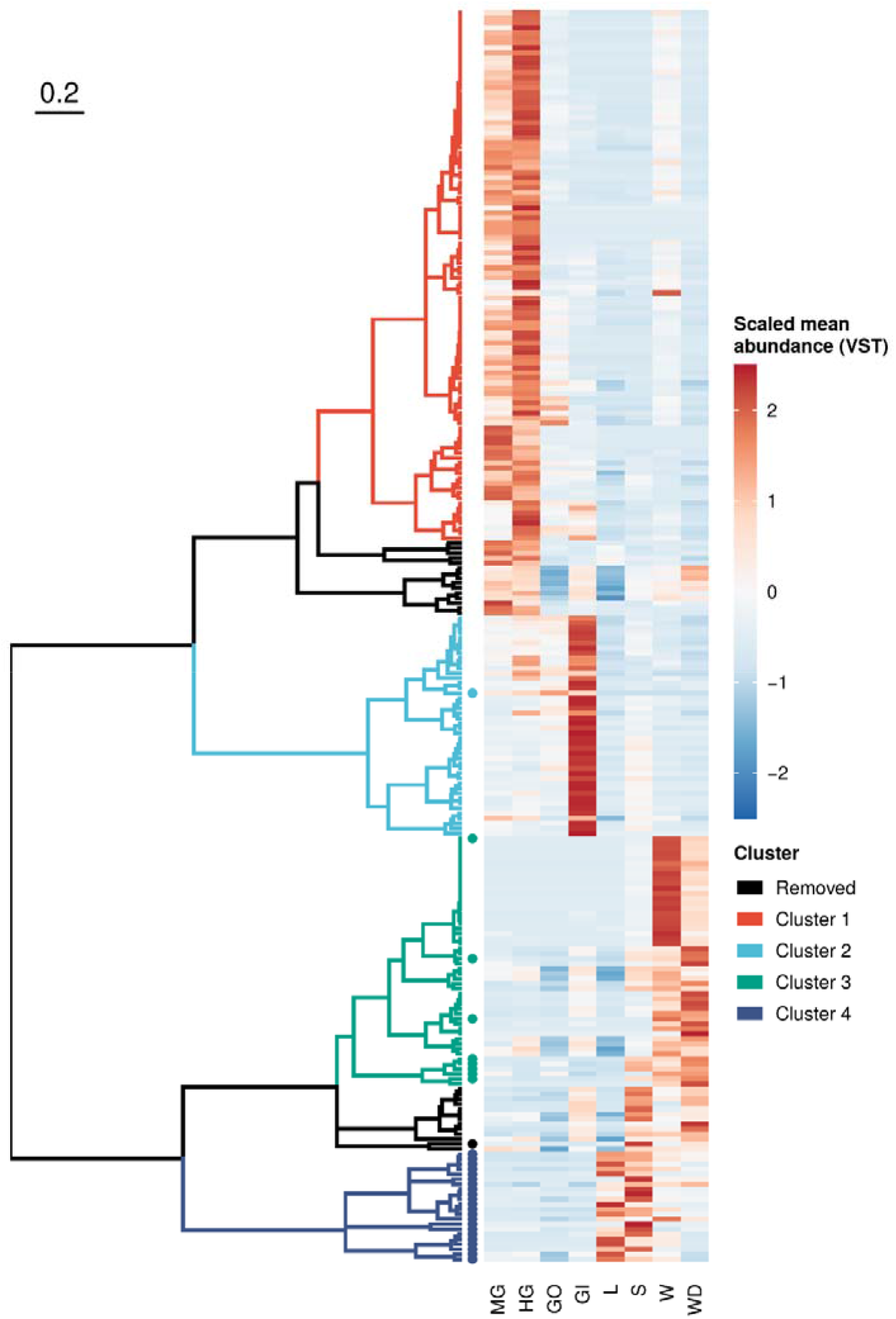
Sequence variant clustering according to their abundance along sample types. Amplicon sequence variants reporting a different abundance pattern in one or more sample types (loglikelihood ratio test of DESeq2) were clustered according to their mean variance-stabilized abundance. Abundance values were reported using different colours after clustering based on Kendall correlation (right side of the plot). Clusters were coloured according to the scheme reported in the legend whereas removed clusters (namely those composed with less than 10 variants) where reported in black. Sequence variants inferred from ITS-1 amplicon sequencing were reported using a solid dot.

### Taxonomic and functional enrichments in crab’s organs and environmental samples

An exploration of the distribution of the scaled variance-stabilized counts within each sample group is shown in Figure 5a. Pairwise Wilkoxon test revealed that the ASVs present in the cluster 1 were significant enriched in the gut, while ASVs included in cluster 2 were significantly enriched in the gills (Figure 5a). Clusters 3 and 4 were significantly related to environmental matrices, cluster 3 to the water and water debris samples, while cluster 4 with litter and soil samples (Figure 5a).

**Figure 5:**
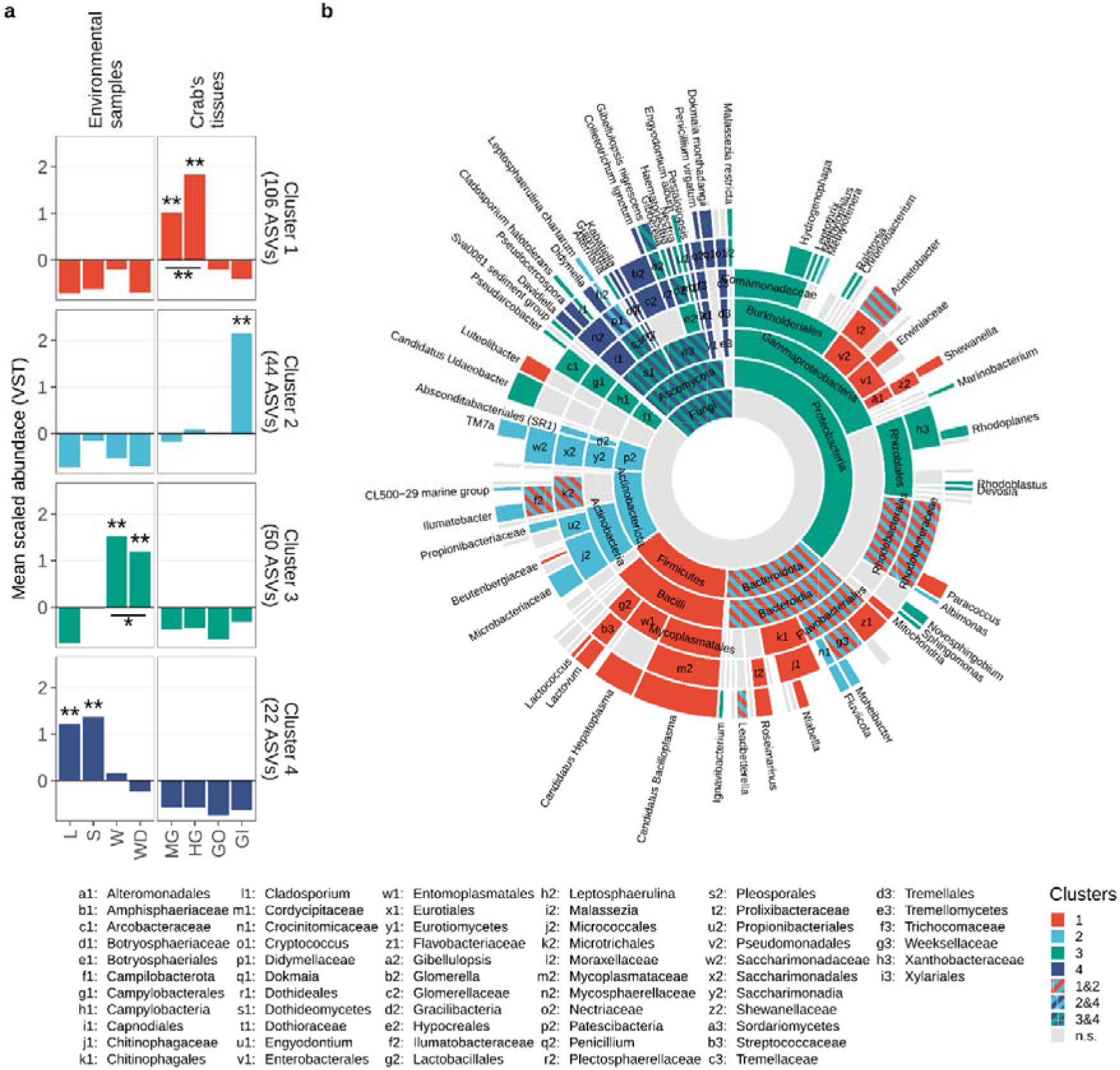
Taxonomic enrichment in clusters. a) Scaled variance-stabilized counts were tested for differences along sample types within all clusters detected. A pairwise Wilcoxon test was performed, and results were reported highlighting significant differences with one asterisk (p-value < 0.05) or two (p-value < 0.01). A complete overview of significant differences was reported in Figure S7. b) Sunburst plot of taxa showing a significant enrichment in a given cluster were coloured according to the colour scale showed in panel a. If a given taxonomy was significantly enriched in more than one cluster, the corresponding sector of the plot was coloured using a striped pattern (as reported in the legend). Leaves represent the most specific level at which a given variant has been classified, namely genus level for 16S rRNA amplicons (Bacteria and Archaea) and species level for ITS-1 region amplicons (Fungi).

Enrichment analysis estimated using log_2_ fold-changes allowed to assign a fair number of ASVs to a specific taxon present in the previously described clusters (Figure 5). Proteobacteria was the most represented phylum of the whole 16S dataset and comprised ASVs belonging to clusters 1, 2 and 3 (Figure 5). The results of this analysis are displayed in the sunburst plot of Figure 5b, which shows that some taxa were enriched in one or more clusters. Significantly enriched taxa were reported from the highest taxonomic rank considered (namely the domain level, the centre of the plot) to the lowest available taxonomic level (namely genus, for bacteria/archaea and species for fungi) and were hierarchically ordered. The intestinal cluster (i.e., cluster 1) was enriched in members of the Firmicutes phylum, more specifically members of Bacilli class including *Candidatus Hepatoplasma, Candidatus Bacilloplasma, Lactovum* and *Lactococcus* (Figure 5b). Members of the phylum Bacteroidota and some taxa related to Protebacteria and Actinobacteriota were enriched in both clusters 1 and 2. Within the phylum Bacteroidota, genera *Roseimarinus* and *Niabella* were associated with cluster 1, *Fluviicola* and *Moheibacter* were related to cluster 2, and *Leadbetterella* was shared between both clusters (Figure 5b). Other clear associations were highlighted, with cluster 3 mainly associated with Proteobacteria and Ascomycota, while cluster 2 mainly related to Actinobacteriota. The genera of these latter phyla are not exclusively associated to a single cluster and are present in multiple clusters. Cluster 4 showed a taxonomic assignment totally related to the Fungi kingdom, in particular to the phylum Ascomycota.

Molecular functions inferred from 16S rRNA gene amplicons showed that bacterial structures associated with crab’s organs are not only taxonomically defined but also functionally defined (Figure 6a). Clusters of bacterial variants were mainly enriched/depleted by a unique set of molecular functions that were absent in other clusters. The inferred genomic content of ASVs detected in crab’s gut (Cluster 1) was enriched by 359 GO terms and depleted by 21, with more than a half (218 terms, 18 depleted and 200 enriched, Table S8) peculiar only to this cluster (Figure 6b). Enzymes involved in the hydrolysis of complex carbohydrates—such as cellulases and xylanases—and commonly associated to the gut of model herbivorous arthropods were not enriched in the gut microbiome of the crab (Table S9). The gills (Cluster 2) had a population of bacteria enriched by 231 GO terms and depleted by 14 terms, with roughly one third (88 terms, 12 depleted and 76 enriched) significantly found only in these organs (Figure 6b). Functions associated to biofilm formation such as cellulose synthase *(30, 31)* were enriched only in this cluster (GO:0016760), whereas functions associated to the nitrogen cycle—such as nitric- and nitrous-oxide reductase activity—were shared between the gills and the environmental cluster 3. The latter, mainly composed of ASVs mostly found in water and water debris samples, had the largest set of GO terms (401 terms, 22 depleted and 379 enriched) with more than a half (223 terms, 19 depleted and 204 enriched) exclusively present in the inferred genome of ASVs highly abundant in that cluster (Figure 6b). Only 4.87% of the total GO terms was significantly enriched in all clusters (37 terms out of 759, with no terms significantly depleted), indicating a functional role of microbial communities both in crab’s organs and in environmental samples (Figure 6b and c, and Table S8). ASVs enriched in crab’s gut and gills shared more functions (GO terms) with the environmental cluster 3 (water and water debris) than between themselves (Figure 6b). In particular, 73 and 68 functions (with only 2 and 1 depleted terms) were shared between the gut and the gills, respectively, and the ASVs detected in the environmental cluster 3 (Figure 6b, Table S8). Considering the total number of terms significantly enriched/depleted, the gills were the organs more impacted by the environment with roughly one third (27.8%) of molecular functions shared with cluster 3. In contrast, the more populated organ, the gut, shared only 19.2% of the total number of functions with the environmental cluster.

**Figure 6:**
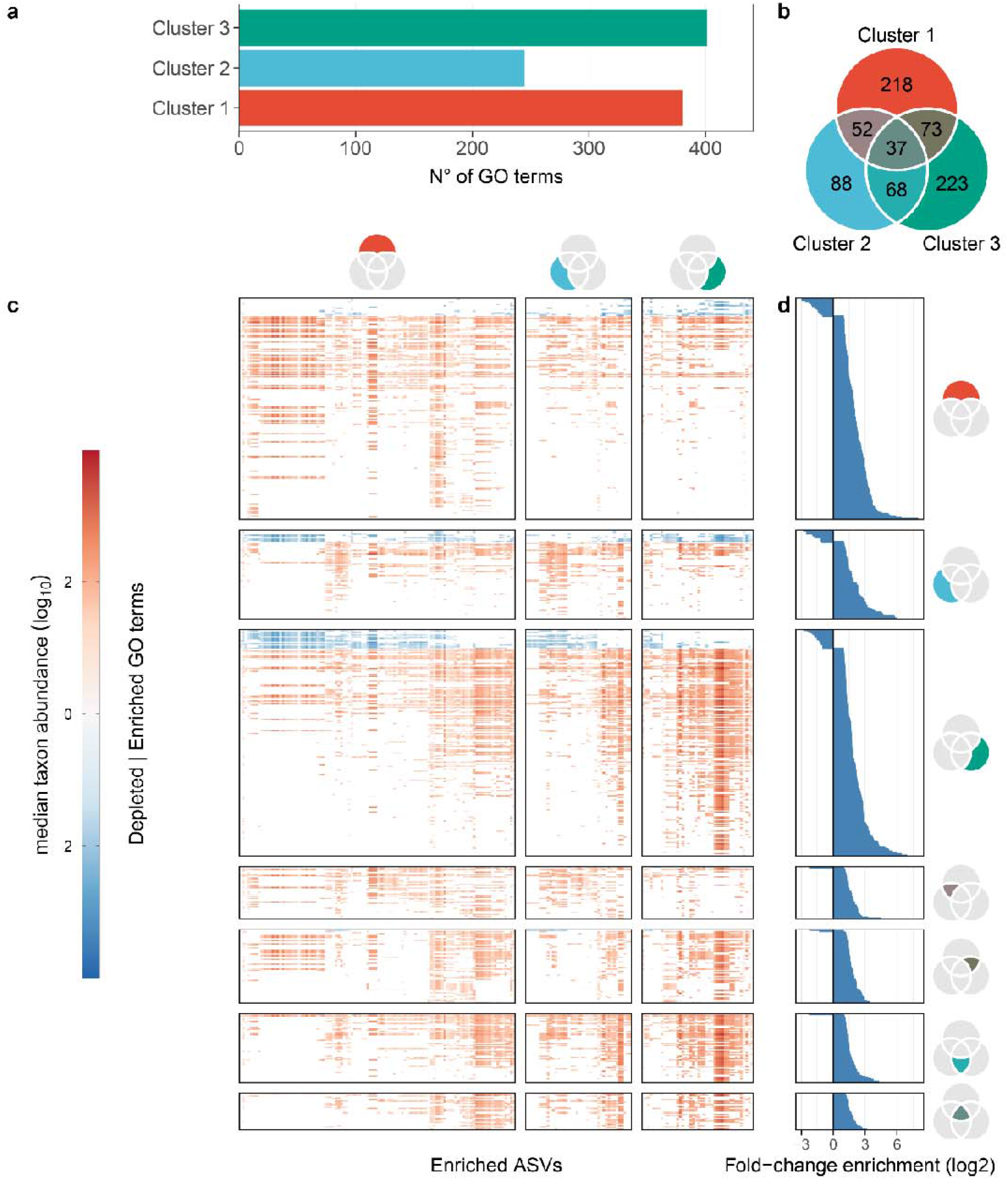
Enrichment analysis of “Gene Ontology” terms associated with detected? functions. **a)** Number of GO terms associated to bacterial function detected in each cluster (cluster 4 was not reported since it was entirely composed of Fungi). b) Venn diagram of GO terms detected in the ASVs of each cluster. Diagram sections in common between two or more clusters were coloured by interpolating colours reported in panel a. c) Median taxon abundance of GO terms (y-axis) in all ASVs detected in each cluster (x-axis). The median taxon abundance was reported using different shades of red—for enriched terms, namely those detected with a higher frequency in respect to the whole population—and blue—for depleted terms, namely those detected with a lower frequency than the rest of the population. The plot was vertically divided according to ASV clusters whereas it was horizontally divided according to venn diagram sections reported in panel b. d) Mean enrichment fold changes associated to each term. Fold-changes were transformed using the logarithmic function (with base equal two) to report enriched and depleted terms symmetrically around the zero.

## Discussion

Our results show the presence of a specific microbiome associate to the semi-terrestrial brachyuran crab *C. haematocheir*, strikingly different from that of the surrounding environment This difference is more pronounced for Bacteria than Fungi. Further, the crab microbiome profiling reveals the presence of consistent organ-specific microbial communities. *C. haematocheir* represents one of the most recent attempts of land colonisation by an arthropod. Consequently, the presence of a specific microbiota suggests its possible role in the host’s adaptation to terrestrial environment. The degree of specificity for both the taxonomic and functional attributes in different organs can be explained by the adaptive evolution of symbiotic host-microbe associations, which now represent a single biological unit under selection from the recently colonised terrestrial habitat, i.e., a holobiont *(sensu 32)*. Temporary host-microbiota associations can be created, retained, or lost over time due to the large number of organisms-microbe interactions. Thus, a crucial test needed to verify the holobiont as a single selection unit is to assess the degree of specificity between the microbial community and its host organism. Our sampling design and analytical approach were able to define stable, organ-specific microbiota in both the gut and gills of individuals belonging to different populations, suggesting a tight and persistent association between those microbial communities and *C. haematocheir*. The organ specificity is supported by the observation that such microbiota were consistently different among the different organs but importantly their composition remained similar across sampling sites, where local microbial assemblages proved to be site specific and different from those associated with the crabs.

The most enriched microorganisms in the gut microbiota of our model species are shared with terrestrial isopods, which moved to the land roughly 300 million years ago, and are supposed to play a central role in their adaptation to a diet based on vascular plant tissues *(33)*. One of the general assumptions of the hologenome concept, however, is that both the host genome and the genetic information of the associated microbiome are transferred from one generation to the next. This postulation underlies the critical role of mothers in transferring microorganisms to their offspring *(34)*, a challenging process for our model organism that has highly dispersive planktonic larval stages followed by a recruitment phase. Another crucial result is that fungal communities are exclusive to the environmental matrices while almost absent in *C. haematocheir* organs. This suggests a specific selection process that favours bacteria versus fungi, possibly supported by selection against yeasts and fungi since several fungi are known pathogens for arthropods and crabs *(21, 35)*. To the best of our knowledge, the present integrated set of results support the idea that semi-terrestrial crabs, only recently migrated onto the land, are associated with microbial communities specifically selected in response to the dramatic environmental pressures posed by the sea-to-land transition. The next questions are where these microbes were acquired and where are their reservoirs?

In the last decades there has been a proliferation of experimental studies supporting the view of holobionts (hosts and their associated microbiota) as units of selection *(36)*. These studies showed how associated microbes have a central role in the host biology, ecology, and evolution, since microbiota are genetically more dynamic and can change more rapidly than the host genome in response to environmental pressures *(32, 37)*. Undeniably, stable symbiotic insect-microbe interactions have been extensively investigated and proved to be crucial for the evolution of many of the several lifestyles of insects *(38, 39)*. Studies on insects, however, cannot shed light on the importance of such associations for the first steps towards land colonisation, which was carried out by insects as far as the Ordovician *(4)*. To bridge this knowledge gap, in this study we focused on a semi-terrestrial brachyuran crab that, from a paleontological point of view, just recently had to face the dramatic challenges represented by excretion, breathing and digestion in terrestrial habitats (for a review see *8)*.

The digestive system of brachyuran crabs comprises different tracts according to their embryological origin and functional role. We intentionally focused on the midgut -where central digestion and absorption occur- and the hindgut -which has a role in water and ion transport *(8, 17)*. In contrast to the intestinal bacterial communities found in the freshwater Chinese mitten crab, *Eriocheir sinensis (40)*, and notwithstanding the different functions, the selected intestinal tracts of *C. haematocheir* were homogeneous in terms of microbial communities, suggesting, at least for this terrestrial crab species, a similar role of such communities.

Host-associated microbiota are known to play a crucial role in the digestive process of many terrestrial arthropods *(37)*. Thus, it is not surprising that the highly diverse microbial community found in *C. haematocheir* gut is clearly distinguished from all the other internal and environmental assemblages. This semi-terrestrial species mainly relies on leaf litter and supplements its diet with small arthropod preys *(28, 29)*. The ability to cope with difficult-to-digest vascular plant compounds (i.e., cellulose, lignin, and polyphenols) has been brought forward as a fundamental trait in the adaptive processes related to terrestrialisation *(17, 41, 42)*. The termites’ ability to degrade lignocellulose is classical example of high gut microbiome specialization *(37)*. Conversely to the termites’ gut microbiome, however, the bacterial assemblages inhabiting the gut of *C. haematocheir* were not enriched in known functions that would help the host to selectively digest cellulose products. Indeed, no gene with cellulase activity were enriched in the gut, possibly reflecting a more variable diet of these crabs, not strictly specialized on plant material digestion *(28, 29)*. The gut microbiome shared limited similarities with the gut core microbiome of a marine predatory brachyuran, the commercially important mud crab *Scylla paramamosain (43)*. Core Proteobacteria shared by *S. paramamosain* and *C. haematocheir* include only *Shewanella*, while Firmicutes include *Lactococcus, Candidatus Hepatoplasma* and *Candidatus Bacilloplasma* (hereafter *Hepatoplasma* and *Bacilloplasma)*. *Hepatoplasma*, and *Bacilloplasma* are rare in marine environments, while they were previously described as colonizers of the hepatopancreas of the terrestrial isopod *Porcellio scaber (33, 44)*. *Hepatoplasma* was detected in *P. scaber* specimens collected from very different geographical areas (Germany and western Canada), proving the strong host-microbe association *(44, 45)*., Other bacterial genera found in our intestinal cluster (such as *Niabella, Paracoccus* and *Shewanella)* were also isolated from terrestrial isopods *(41)*. Of particular interest is also the presence of *Lactococcus*, which is a genus of lactic acid bacteria known as homofermenters, producing a single product, lactic acid, as the main, or only product of glucose fermentation. Lactic acid has a selective role since it lowers the pH and selects the environmental microbes that can potentially thrive in the gut. These microorganisms, common in the dairy industry, are also known to colonize the gut of termites, *Lactococcus nasutitermitis (46)* and do not include marine species, although they are occasionally described as pathogens of aquacultured species *(47, 48)*. Another microorganism we found in the intestinal cluster that has never been described in the marine environment, thus suggesting acquisition from terrestrial microorganisms, is *Lactovum*, a genus of bacteria within the family Streptococcaceae. The genus includes a single species, *Lactovum miscens*, an aerotolerant, anaerobic species originally isolated from soil of the Stiegerwald forest in Germany *(49)*. In addition, the intestinal cluster was enriched in *Erwiniaceae*, a family that includes insect’s symbionts such as *Buchnera aphidicola (50)*. The gut microbiota we found in the mid- and hindgut of *C. haematocheir* then shared similarities with the gut microbiota of other terrestrial crustaceans and insects, suggesting the presence of functions/preferences of those taxa that could help arthropods digestive systems, a hypothesis that deserves further functional investigations.

The possibilities of a vertical transmission vs. acquisition from the environment of this specialized gut microbiota deserves a deeper discussion and further experiments. First, land and terrestrial crabs are neither social or gregarious and only few of them show some basic degrees of parental care *(18, 51)*. Second, *C. haematocheir*, as well as the most land-adapted family of brachyuran crabs, the Gecarcinidae, still retains an indirect development strategy and releases planktonic larvae in coastal waters. Consequently, *C. haematoicher*, as well as the land crabs with planktonic larvae in general, may have developed intimate gut-specific symbiotic associations only through pseudo-vertical transmission (sensu *53)*, i.e., by acquiring and selecting their gut microbiota directly from food or, most probably, through consumption of adults’ faeces (coprophagy) found in or around their burrows. The megalopae of various terrestrial and land crabs are known to specifically recruit at their spawning grounds, as showed by the iconic mass recruitment of the Christmas Island red crab *Gecarcoidea natalis (53)*, or in areas where they can chemically detect adult populations, such as the case of the swamp ghost crab *Ucides cordatus (54)*. In some species, this selective recruitment towards populations of conspecific adults ultimately results in the occurrence of juveniles and sub-adults within secondary branches excavated along the burrows of adult crabs *(55–57)*. Indeed, these behaviours maximise the chances of finding adult populations and, consequently, a successful pseudo-vertical transmission of microbiota through coprophagy. Further metagenomic analyses on faeces are needed to clarify this point.

The gonads host a low microbial diversity with respect to the other organs and environmental matrices we analysed, as expected for an internal organ in no direct contact with the environment and not morphologically connected to the gut and the gills. Their microbial community, moreover, is shared with both the other organs and the environment, determining the absence of a specific taxonomic and functional cluster associated with them. Microorganisms are known to be associated with the reproductive system of arthropods, in both insects *(58)* and crustaceans *(59)*. Few studies, however, have specifically characterised these gonad-associated microbiota and most of them were focused on specific pathogens of sexually transmitted infections or reproductive parasites, with the most notable example being *Wolbachia (60)*. When present, the vertical transmission of specific microbiota during oogenesis or at birth is known to determine a colonization of the gonads that ultimately affects the microbial composition of the offspring *(59)*. In this view, the absence of an exclusive microbiota in *C. haematoicher* gonads can be explained, as for the gut microbiota, by the lack of vertical transmission through the parental-offspring pathways, because of the presence of planktonic stages.

Although protected in the gill chambers, *C. haematocheir* gills are in direct communication with the external environment, but they do show a unique resident microbiota, which is different from the microbial communities detected in the environmental matrices. These gill-associated microbial assemblages are only enriched in prokaryotes, unlike the soil/litter and the water communities, which are composed almost entirely by fungi and a mix of Proteobacteria and fungi, respectively. The prokaryotic cluster associated with the gills is characterised by a strong uniformity in terms of both taxonomy and functions. Actinobacteria associated to the gills include Microbacteriacea and *Illumatobacter*, which are known to play an important role in marine organisms by producing bioactive compounds crucial in the defence against pathogens *(61)*. Some of the bacteria detected on the gills of *C. haematocheir*, such as *Ilumatobacter* and *Albimonas*, were also found in the gills of the Chinese mitten crab *Eriocheir sinensis (40)*, but at this stage their metabolic functions are not clear. In contrast to the gut, genes associated with cellulose synthase activity were found in the microbiota associated with these organs (GO:0016760). This activity is typical of biofilm-forming bacteria that use cellulose both as a physical barrier against armful molecules such as antibiotics but also biocides and metallic cations, and as a molecular glue to help their interaction with the host *(31)*. Since the gills of *C. haematocheir* are exposed to external perturbations, the presence of biofilm-forming functions may help to boost the resilience of bacteria stabilizing host-microbiome interactions. Genes related to the reduction of nitric compounds (namely: nitric oxide reductase activity, GO:0016966, and nitrous-oxide reductase activity, GO:0050304) were enriched in both the gills and environmental water and water debris samples. Besides their role in the anaerobic metabolism of nitrogen, these functions are involved into pathogenesis and antibiotic resistance in bacteria *(62)* and may help tissue colonization. In addition, the covalent incorporation of a nitric oxide molecule (nitrosylation) of cytochrome c and quinol oxidases inhibits cellular respiration acting as an antimicrobial molecule. We speculate that this mechanism may be used by bacteria inhabiting the gills of *C. haematocheir* as a possible molecular defence against external pathogens helping the crab, and themselves, to thrive in different terrestrial environments.

Our results show that fungi are strongly depleted in all crab’s organs and that gills should be considered an efficient selective filter between the environment and the host. In our opinion, the substantial absence of fungal associations can be explained by different hypotheses that all converge on the existence of defences from potential pathogens. We may hypothesize that some physiological and anatomical characteristics of specific organs, such as gut and gills, may inhibit adhesion and development of fungi. In insects, for instance, physiochemical characteristics of the cuticle, or feeding habits, can counteract the entomopathogenic fungi cuticular adhesion, germination of spores and hyphal growth *(63)*. Moreover, bacterial production of antifungal molecules may limit the growth of fungi. Events of fungal exclusion by direct competition mediated by symbiotic bacteria have been extensively described in beetles *(64)*. *Streptomyces* sp. SPB74, symbiont of the beetle *Dendroctonus frontalis*, produces an active compound, mycangimycin, which specifically inhibits antagonist fungi without acting on the mutual ones *(64)*. This protective role may be extended to the offspring. Antifungal molecules produced by the symbiotic bacterium *Burkholderia gladioli* are known to protect the eggs of the beetle *Lagria villosa* against pathogenic fungi *(35, 65)*. Pathogenic fungi have been reported in intertidal and terrestrial crabs, such as the case of the swamp ghost crab, *U. cordatus*, infected by species of black yeast, *Exophiala cancerae*, and *Fonsecaea brasiliensis*, which cause a condition called “lethargic crab disease” *(21, 22)*. The role of prokaryotic associations in the defence against pathogenic fungi can be of critical importance for terrestrial crabs, since the immune molecules of crustaceans are less efficient than those of insects against fungal infections *(66)*.

In conclusion, this is the first attempt to ascertain the role of host-microbiome associations in the transition to land of brachyuran crabs, the most recent arthropod taxa to perform such an evolutionary leap. We found clear differentiations among the microbiota associated with different crab’s organs and the ones found in the environmental matrices, suggesting a selective acquisition of the microbiota through “gain” and “loss” mechanisms. The differences found among the organ-related and environmental clusters were not merely taxonomical since the different clusters harbour different metabolic profiles. These results corroborate the hypothesis that the recorded differences can be due to the presence of metabolic complementation mechanisms that took place in those organs mostly impacted by the challenges posed by the terrestrial environment. Our data also show possible evolutionary convergences towards a uniform ‘terrestrial intestinal microbiota’ across different lineages of arthropods, suggesting the presence of bacterial associations linked to terrestrial life. The present study demonstrates mechanisms of specific microbial selection in the sesarmid crab *C. heamatocheir* and strongly supports the hypothesis that semi- and terrestrial crabs are an appropriate model system to study the evolution of arthropod-microbe interaction under the selective pressures posed by the sea-land transition, which is happing right now in Brachyura *(23)*.

## Material and methods

### Study species

*Chiromantes haematocheir* (Decapoda; Brachyura; Sesarmidae) is a semi-terrestrial crab colonising the coastal vegetated areas from Taiwan to South East Asia *(28, 67)*. In Hong Kong, it forms large populations in areas of lowland secondary forest adjacent to mangroves and in pockets of riverine forests, where it was also observed climbing trees. Very little is known about its ecology, apart from the fact that it digs deep burrows and, as in many sesarmids, it releases pelagic larvae into the ocean. This species was selected because it is the most land adapted among the Hong Kong brachyuran crabs.

### Sample collection and total DNA extraction

We selected three large populations of *C. haematocheir* that colonised distant catchments across the Hong Kong territory. The selected populations were sampled at Shui Hau (Southern Lantau Island), To Kwa Peng (Eastern coast of Sai Kung Country Park, New Territories), and Sai Keng (Three Fathoms Cove, Tolo Harbour New Territories). From each site, eleven sexual mature adult crabs (carapace width range between 13.4 mm and 32.2 mm) were collected in October 2018. To explore differences in microbial community composition across the various crab’s organs and the surrounding environment, we collected and analysed four organ samples (gills, hindgut, midgut and gonads) and four environmental matrices (sediment, leaf litter, freshwater and freshwater debris). Due to the intensive sampling, all environmental samples and alive crabs were immediately frozen and subsequently transported to the laboratories of the Division of Ecology and Biodiversity (The University of Hong Kong). The dissections were then performed under sterile conditions. All dissection instruments were sterilized over an open flame to eliminate residual DNA and washed with 75% EtOH to prevent cross-contamination. After removing the carapace, gills, gonads, hindgut and midgut from each crab were excised under a stereomicroscope and stored at -20°C in RNAlater (Thermo Fisher Scientific) stabilization solution until DNA extraction.

Total DNA extraction from crab organs, sediment, leaf litter and freshwater debris was performed using the DNeasy PowerLyzer PowerSoil Kit (QIAGEN) following manufacturer’s protocol. Total DNA extraction from water samples was performed using the DNeasy PowerWater Kit (QIAGEN) following manufacturer’s protocol, after having filtered 100ml of freshwater through 0.2μm Thermo Scientific™ Nalgene™ Sterile Analytical Filter Units (Thermo Fisher Scientific). Extracted DNA samples were stored at -20°C. Before the DNA library’s preparation, DNAs were quantified fluorometrically by using Qubit dsDNA HS Assay Kit (Thermo Fisher Scientific).

### 16S (V3-V4) rRNA gene amplification and sequencing

The preparation and sequencing of the 16S library were performed at Laboratory of Advanced Genomics, Department of Biology, University of Florence (Firenze, Italy). PCR amplifications of the bacterial V3-V4 16S rRNA gene fragments were performed using KAPA HiFi HotStart ReadyMix (Roche) and the primer pair 341F (5’-CCTACGGGNGGCWGCAG-3’) and 805R (5’-GACTACNVGGGTWTCTAATCC-3’) *(68)* with overhang Illumina adapters. 16S amplicon PCR protocol was set on 25 μl of final volume. In detail for each reaction, 12.5 μl of 2× KAPA HiFi HotStart ReadyMix (Roche), 10 μl of 1μM forward and reverse primers and 2.5 μl of template DNA (5–20 ng/μl) were combined. Amplicon PCR reaction was performed using the GeneAmp PCR System 2700 (Thermo Fisher Scientific) and the following cycling conditions: denaturation step at 95 °C for 3 minutes; 35 (gonads, water and water debris) and 25 (hindgut, midgut, gills, soil and litter) cycles: at 95 °C for 30 seconds, 55 °C for 30 seconds, 72 °C for 30 seconds; final extension step at 72 °C for 5 minutes. All PCR products were checked through electrophoresis on 1.5% agarose gel and then purified using KAPA Pure Beads (Roche) following the manufacturer’s instructions. To apply the Illumina adapters sequencing indexing using Nextera XT Index Kit V2 (Illumina), a second PCR amplification was then performed by preparing a reaction mix in accordance with the 16S metagenomic library preparation protocol *(69)*. An indexing step was made for all samples by seven PCR cycles. Amplicon products from indexing PCR were purified using KAPA Pure Beads (Roche) and their quality check was performed using Agilent 2100 Bioanalyzer (Agilent Technologies) with Agilent DNA 1000 Kit (Agilent Technologies). Subsequently, concentration check was performed by Qubit dsDNA HS Assay Kit (Thermo Fisher Scientific). Finally, the barcoded libraries were balanced and pooled at equimolar concentration, before being sequenced on an Illumina MiSeq (PE300) platform (MiSeq Control Software 2.6.2.1).

### ITS1 rDNA region sequencing

ITS1 library preparation and sequencing were performed at the Research and Innovation Centre, Fondazione Edmund Mach (FEM) (S. Michele all’Adige, Trento, Italy). Fungal ITS1 fragments were amplified by PCR using the FastStart High Fidelity PCR System (Roche) for environment matrixes and the Hot Start High-Fidelity DNA Polymerase (NEB) for animal matrixes following the manufacturer instructions using the primers ITS1F (5’-CTTGGTCATTTAGAGGAAGTAA-3’) *(70)* and ITS2 (5’-GCTGCGTTCTTCATCGATGC-3’) *(71)* with overhang Illumina adapters. ITS1 PCRs were performed in 25 μl of final volume. In detail for each reaction, 2ul of 10uM forward and reverse primers were used in combination with 1 µl of template DNA (5-20 ng/ul). Amplicon PCR reaction was performed using the GeneAmp PCR System 9700 (Thermo Fisher Scientific) and the following cycling conditions: denaturation step at 95 °C for 3 minutes; 25 (litter) and 33 (all other samples) cycles: at 95 °C for 20 seconds, 50 °C for 45 seconds, 72 °C for 90 seconds; final extension step at 72 °C for 10 minutes. All PCR products were checked on 1.5% agarose gel and purified using the CleanNGS kit (CleanNA, the Netherlands) following the manufacturer’s instructions. Subsequently a second PCR was performed to apply the Illumina sequencing adapters Nextera XT Index Primer (Illumina) *(69)*. The indexing step was made for all samples by seven PCR cycles. After Indexing PCR amplicon libraries were purified using the CleanNGS kit (CleanNA, the Netherlands), and the quality control was performed on a Typestation 2200 platform (Agilent Technologies, Santa Clara, CA, USA). Afterwards all barcoded libraries were mixed at equimolar concentration, quantified by qPCR Kapa Library quantification kit (Roche) and sequenced on an Illumina MiSeq (PE300) platform (MiSeq Control Software 2.5.0.5 and Real-Time Analysis software 1.18.54.0).

### Amplicon sequence variant inference

The DADA2 pipeline version 1.14.1 *(72)* was used to infer amplicon sequence variants (ASVs) from raw sequences. Primers used for PCR amplification were removed using cutadapt version 1.15 *(73)* in paired-end mode. If a primer was not found, the sequence was discarded together with its mate to reduce possible contamination. For ITS amplicon sequences reads containing both the forward and reverse primers were considered valid only if concordant (one of the two primers must be present but reverse and complemented) with a cut-off length of 70bp. Low quality reads were discarded using the “filterAndTrim” function with an expected error threshold of 2 for both forward and reverse read pairs (namely only reads with more than 2 expected errors were removed). Denoising was performed using the “dada” function after error rate modelling (“learnErrors” function). Denoised reads were merged discarding those with any mismatches and/or an overlap length shorter than 20bp (“mergePairs” function). Chimeric sequences were removed using the “removeBimeraDenovo” function whereas taxonomical classification was performed using DECIPHER package version 2.14.0 against the latest version of the pre-formatted Silva small-subunit reference database *(74)* (SSU version 138 available at: http://www2.decipher.codes/Downloads.html) and the Warcup database for fungal ITS1 *(75)*. All variants not classified as Bacteria, Archaea or Fungi were removed together with sequences classified as chloroplasts or mitochondria (16S rRNA sequences only). Additional information on the sequence variant inference pipeline used were reported in Supplementary Materials.

### Inferring functional content of amplicon variants

The genome content of bacterial ASVs was inferred using PICRUSt2 pipeline *(76)*. Enzyme Commission Numbers (EC numbers) were converted into Gene Ontology terms (GO terms) using the mapping file available at: http://www.geneontology.org/external2go/ec2go. Gene abundance was retrieved using the “--strtified” option to report gene abundances at species level (ASVs. Additional information about functional content inference was reported in Supplementary Material.

### Statistical analyses

All statistical analyses were performed in the R environment (version 3.6). Briefly, alpha- and beta-diversity analyses were conducted using the vegan package version 2.5 *(77)* in combination with the iNEXT package version 2.0 *(78)*. Normalization and differential abundance analyses were performed with DESeq2 version 1.28 *(79)* whereas enrichment analysis of taxa and functions was performed using hypergeometric test (“phyper” function of R stats package). For additional details about tests data manipulation see Supplementary Materials.

## Acknowledgments

We thank the friends of the Integrate Mangrove Ecology laboratory of the Swire Institute of Marine Science, HKU for their help in sampling and lab work. We also thank Prof Colin Little and Gray A. Williams for inspiring and discussing the original idea behind the study.

## Author contributions

G.B. performed bioinformatics analyses and wrote the manuscript; S.F. supervised lab work and wrote the manuscript; N.M. performed 16S laboratory analyses and wrote the manuscript; C.L.Y.C. and K.H.N carried out the samplings and performed DNA extractions; D.C. and A.M. helped conceiving the original idea; and S.C. conceived the original idea, collected the samples and wrote the manuscript. S.C. and D.C provided funding for the experiments. All the authors commented and validated the manuscript, participated to plan the sampling design and critically discussed the results.

## Data and materials availability

data needed to evaluate the conclusions are present in the paper and in Supplementary Materials. All codes used in the work have been uploaded to a public Github repository available at: https://github.com/GiBacci/Chiromantes_haematocheir_microbiome. Sequencing data were uploaded to the European Nucleotides Archive (ENA) under the project ID: PRJEB43930.

