## Supplementary materials for "Conserved organ-specific microbial assemblages in different populations of a terrestrial crab"

Supplementary material
Organ-specific microbiota enhances the terrestrial lifestyle of a brachyuran crab

Giovanni Bacci^1^*, Sara Fratini^1^*, Niccolò Meriggi^1^*, Christine L. Y. Cheng^2^, Ka Hei Ng^2^, Massimo Pindo^3^, Alessio Iannucci^1^, Alessio Mengoni^1^, Duccio Cavalieri^1^†, Stefano Cannicci^1,2^†

^1^ Department of Biology, University of Florence, Sesto Fiorentino, IT50019, Italy
^2^ The Swire Institute of Marine Science and the Division of Ecology and Biodiversity, The University of Hong Kong, Hong Kong, Hong Kong SA
^3^ Research and Innovation Centre, Fondazione Edmund Mach, San Michele all’Adige, IT38098, Italy

* These authors contributed equally to this work

#### Sequence variants inference

Amplicon sequences obtained from 16S rRNA gene and ITS1 region were analysed using the DADA2 pipeline (version 1.10.1) (*1*) in R version 3.5.3 (<http://www.R-project.org>). Primers used for PCR amplification were removed using cutadapt version 1.15 in paired-end mode (*2*). Sequences that did not contain the primer were discarded removing both pairs if at least one of them (R1 or R2) did not pass the filtering criteria (96.26% and 90.58% of raw sequences coming from 16S rRNA gene and ITS1 region were retained, respectively). Only the corresponding primer was matched in bacterial 16S rRNA amplicons (341F for R1 mates and 805R for R2 mates) whereas in ITS1 regions both primers could be matched but only if identified in the correct orientation (ITS1F forward in 5’ position and ITS2 reverse and complement in 3’ position for R1 mates and the opposite for R2 mates; for additional information see <https://benjjneb.github.io/dada2/ITS_workflow.html>). Since samples were sequenced using multiple Illumina flow cells, we followed the “big data” approach (described at: <https://benjjneb.github.io/dada2/bigdata.html>) to infer amplicon sequence variants (ASVs) with a relatively modest memory requirement and to respect the dependence of DADA2 error models from the sequencing run. Reads coming from 16S rRNA gene amplification (V3-V4 region) were trimmed at 280bp (forward) and 200bp (reverse) to remove low quality segments while retaining enough overlapping nucleotides to reconstruct the full amplicon (*3*, *4*). Fungal ITS1 amplicons were not trimmed to a fixed length to avoid losses of longer fragments that could be present due to the high variability in length of this region, as also reported by the authors of the DADA2 pipeline (<https://benjjneb.github.io/dada2/ITS_workflow.html>). On the other side, to reduce the number of small fragments, fungal reads shorter than 70bp were removed from subsequent analyses. Reads (and their respective mate) that contained ambiguous bases or more than two expected errors were filtered out to minimize errors due to the sequencing process. Sequences were denoised using the DADA2 algorithm with default parameters. The denoised mates were merged and all reads with any mismatches and an overlap length shorter than 20bp were removed (87.18% and 96.65% of trimmed reads coming from 16S rRNA gene and ITS1 region were correctly merged, respectively). Chimeric sequences were identified and removed using the removeBimeraDenovo function with “consensus” method (5.51% and 0.89% of analysed sequence coming from 16S rRNA gene and ITS1 region were flagged as chimeric and removed from subsequent analyses). Samples with a low coverage (expressed as the number of reads assigned to an ASV) where re-sequenced and added to the inference pipeline (Table S1). To respect the dependence of DADA2 error models from the run, counts from re-sequenced samples were summed together after the variant inference step.

#### Classification of amplicon sequence variants

Sequence variants were taxonomically annotated using the DECIPHER package (version 2.10.2) against two widely used databases: Silva SSU (version 138) (*5*, *6*) and Warcup v2 ITS database (*7*). The IDTAXA algorithm implemented in the package (*8*) needs a trained database for the correct classification of query sequences so we used the trained version of both databases available from the DECIPHER site (<http://www2.decipher.codes/Downloads.html>). Sequence variants not assigned to Bacterial, Archaea, or Fungal domains were removed from subsequent analyses to minimize biases due to DNA contamination, or to sequencing errors. Even if the primers used in this work are not reported to cross-amplify chloroplast or mitochondrial DNA (*9*–*11*) we removed any 16S rRNA amplicon sequence variant (ASV) that was assigned to these orders to avoid additional biases during downstream analyses. Two fungal samples, namely samples “GI_TKP_5” and sample “MG_SH_1”, were removed from downstream analyses since their ASV count dropped to zero. The number of reads retained after each step was reported in Figure S1.

#### Fungal amplicon sequence variant evaluation

Fungal variants showed a massive loss of reads in crab’s organs (Figure S1) so we sequenced two samples for each crab’s organ independently in order to validate the workflow used in an unsupervised fashion (Table S1). DNA was extracted and amplified independently for each replicate before sequencing. Replicates were included in the DADA2 pipeline together with the other samples and they were used to evaluate the overall accuracy. We took into account two main parameters: a) the Spearman’s rank correlation coefficient ($\rho$) between replicates, and b) the overall accuracy. The accuracy was computed as:

$$\frac{(TP+TN)}{(TP+TN+FP+FN)}$$

where, $TP$ was the number of ASVs detected in both replicates (true positives); $TN$ was the number of ASVs not detected in both replicates (true negatives); $FP+FN$ was the number of ASVs detected in one replicate but not in the other one (false negatives). Accuracy was estimated by comparing replicates in pairs (two pairs per organ for a total of eight contrasts). We used a non-parametric correlation coefficient to measure monotonic relationships between the abundance of ASVs in all replicates. To reduce differences due to the total size of different replicates, ASV counts were transformed into “parts per thousands” by dividing them by the total read count and then multiplied by 1000. A log_2_ transformation was performed before correlation coefficient calculation (*12*). Values of accuracy and $\rho$ for each sample sequenced were reported in Table S2 whereas the log_2_-transformed counts of ASVs in each replicate was reported in Figure S2.

### Microbial diversity within sample types and sampling sites

The coverage of microbiome data obtained through amplicon sequencing is extremely variable across samples due to differences in PCR amplification, sequencing yields, and clustering algorithms. This variability may induce over/underestimation biases when comparing microbial diversity, especially when using indices that rely on the abundance of single species. To minimize possible biases, we normalized counts by using the “median ratio method” implemented in the counts function of the DESeq2 R package (version 1.28.1) (*13*, *14*). Differently from the “variance stabilizing transformation” (VST)–mainly used within the same package to transform count data into approximately homoskedastic data that can be used for clustering and machine learning approaches–the “median ratio method” produces positive abundance values that can be directly used to estimate differences within- and between-samples (also called alpha- and beta-diversity) (*15*).

Normalized counts were used to estimate differences in the total microbial diversity across sample types by using three alpha-diversity metrics, all based on the inverse Simpson index. This index is defined as the reciprocal of the Simpson index $D$ ($D'$) but it can be also defined as a diversity measure with an Hill number of order two–a definition mostly used in ecological studies (*16*). Hill numbers (${}^{q}D$) are represented with the equation:

$${}^{q}D=\left( \sum_{i=1}^{R} p_{i}^{q} \right)^{1/(1-q)}$$

were $q$ is the order of the Hill number ${}^{q}D$. For $q=2$ we can write the equation as:

$${}^{2}D=\frac{1}{\sum_{i=1}^{R} p_{i}^{2}}$$

Since $\sum_{i=1}^{R} p_{i}^{2}$ corresponds to the definition of the Simpson index, the equation corresponds to the inverse of the Simpson index. This index is particularly useful since it transforms the Simpson concentration–that is by definition the probability of drawing two equal species taken at random from the dataset–into a measure of diversity so that the higher the index, the higher the microbial diversity. We used the iNEXT R package (version 2.0.20) (*17*) to compute interpolated, observed and extrapolated values of alpha diversity with Hill number of order 2 (inverse Simpson index). All values were computed separately for 16S rRNA and ITS1 amplicons by using 30 bootstrap replications each. In order to compare different samples, we choose a fixed range of sizes (number of read sequenced) for the two amplicons so to obtain vectors of diversity of the same size. Since the number of reads collected for 16S rRNA amplicons was higher than that obtained for ITS1 amplicons (a maximum of roughly 460’000 normalized counts for 16S amplicons against 60’000 for ITS1) we computed alpha diversity along 300 and 100 size knots, respectively. The iNEXT package calculate interpolated and extrapolated values of diversity as a function of the sample size (called $m$) using the formula reported in the paper from Chao and colleagues (*18*). In particular an interpolated diversity estimator (${}^{q}\hat{D}(m)$) is derived for any size $m<n$ where $n$ is the total counts in the sample. Similarly, an extrapolated diversity estimator is derived (${}^{q}\hat{D}(m+n)$) for any enlarged sample of size $m+n>n$. Observed microbial diversity is computed by using the diversity formula described above. Confidence intervals were constructed using the bootstrap method and joined to recreate a smooth curve (*19*). Generated curves were extended to the maximum sampling size of all samples in order to check if the sequencing effort could be considered enough for the comparison of alpha-diversity across different samples (namely, the inverse Simpson index). Results were reported for each sample type (Figure 2 panels a and c in the main text) and for each sample site (Figure S3 panels a and c). Average differences between extrapolated and observed inverse Simpson index (Table 1 in the main text) were really low, ranging from 0.05 to 2.74 indicating that the sequencing effort was enough to explore the alpha diversity of sample coming from different biological matrices (both crab’s organs and environmental samples). Observed inverse Simpson index and Good’s coverage values were reported for each sample in Table S3. Good’s coverage index was computed following its standard definition (*20*):

$$(1-\frac{n}{N})\cdot100$$

where, $n$ is the number of sequences found once in a specimen (also called singletons) and $N$ is the total number of counts in that specimen. Good’s coverage estimator ranged between 99.89% and 100.00% across all samples indicating that roughly 0.11% of the reads in a sample came from ASVs that appear only once in that sample. To detect differences across sample types and sampling sites, we used the non-parametric pairwise Wilcoxon test with Benjamini–Hochberg p-value correction. Statistical significance was then reported using alphabetical letters obtained with the multcompLetters function of the multcompView package (version 0.1.8). Results were reported for each sample type (Figure 2 panels b and d in the main text) and for each sample site (Figure S3 panels b and d).

### Microbial diversity between sample types and sampling sites

The variation of microbial community between samples (beta diversity) was assessed using different approaches on microbial counts generated by 16S rRNA and ITS1 amplicon sequencing separately. Before computing any beta diversity index, samples were normalized as explained in the previous section and transformed into relative abundances to smooth differences in coverage across samples. Sample distribution was reported using principal coordinates analysis (PCoA, also called classical multidimensional scaling or MDS) on Bray-Curtis diversity index (cmdscale function of stats package, and vegdist function of vegan package (*21*), respectively). We used the quantitative version of the Bray-Curtis index with formula:

$$d_{jk}=\frac{\sum_{i=1}^{N} |x_{ij}-x_{ik}|}{\sum_{i=1}^{N} x_{ij}+x_{ik}}$$

where $d_{ij}$ is the Bray-Curtis diversity between sample $i$ and $j$, $N$ is the total number of ASVs in the two samples, $x_{ij}$ is the abundance of the ASV $i$ in sample $j$, and $x_{ik}$ is the abundance of the same ASV in sample $k$. Permutational multivariate analysis of variance (PERMANOVA) was used to inspect differences between sample types and sampling sites (adonis2 function of the vegan R package, version 2.5.6) with 1000 permutations and Bray-Curtis diversity index. Sample types and sampling sites were tested in all samples whereas for crab’s organs the crab was added as a covariate for controlling variance due to the crab itself (also called batch effect). The proportion of variance explained by each factor (R squared) was reported in Table S4. Approaches based on permutations, such as PERMANOVA, rely on the assumption of homoscedasticity–the dispersion should be equal in all groups otherwise it is not possible to discern a location effect (the groups) from a dispersion effect (the difference in dispersion). To test this assumption we used the betadisper function (vegan package) to compute distances between each entity (samples) and group centroids. Differences in dispersion were then tested using the analysis of variance (ANOVA) and results were reported in Table S5. Pairwise PERMANOVA was used to assess whether sample types and sampling sites differed based on ASV counts obtained from crab’s organs and environmental samples. The r-squared value of each contrast was used to inspect the amount of variance explained by crab’s organs and environmental samples on the separation between sites and sample types. Principal coordinates analysis was reported in panel a of Figure 3 (by sample type) and Figure S4 (by sampling sites) whereas results of pairwise PERMANOVA were reported in panel b of the same figures.

To detect the number of species present/absent in all sample types and in all their intersections we used a graphical visualization called “upset plot” (upset function of the UpSetR package version 1.4.0 (*22*)). This type of representation is extremely useful to display a large number of intersections that cannot be easily grasped with Venn plots (usually four or more sets)–the number of intersections corresponds to the binomial coefficient which is 15 for four intersections and it roughly doubles passing from four to five (31 intersections). Upset plots were introduced to address this issue by reporting set intersections in a matrix layout where each row is a different set and each intersection–indicated with linked dots–is reported column wise. The size of each intersection is reported with a bar on the top of the corresponding column and is usually ordered according to the degree of the intersection (namely, the number of sets included) and the size (*23*). The number of ASVs shared across different sample types was reported for 16S and ITS1 amplicons (Figure S5) both according to sampling sites (panels a and c) and sample type (panels b and d).

### Clustering of sequence variants

To detect pattern of ASVs across sample types we used the likelihood ratio test (LRT) implemented in the DESeq2 package (version 1.28.1). The main advantage of this approach is that it tests the effect of all levels of one (or more) factor/s at the same time by comparing the likelihood of the full model against a model lacking the factor/s of interest. Since we wanted to identify pattern across different sample types, and given that we had to deal with eight distinct sample types (namely a factor with eight levels), we tested a full model including sampling sites and sample types against a reduced model including only sampling sites. This way we accounted for dispersion due to different sampling sites while testing for the effect of all sample types at the same time. Singletons–amplicon sequence variants detected only once in the whole community–were removed before model fitting to reduce computational time and to prevent ungrounded speculations based on sporadic species. To select the model that best fitted our data, we fitted all the possible fit types implemented in the DESeq2 package (namely, “local”, “mean”, and “parametric”). The choice of the best model was based on the median absolute value of the log-residuals (MALR), defined as:

$$MALR=median(|log(\alpha_{i}^{gw})-log(\alpha_{tr})|)$$

where $\alpha_{i}^{gw}$ is the dispersion estimate of ASV $i$ obtained *a priori* from the data and $\alpha_{tr}$ is the fitted dispersion value obtained from the model (Figure S6, for in-depth definitions of parameters reported see the original DESeq2 paper (*14*)). The model with the lowest MALR value was selected and used for subsequent analyses (fit type “local”). Results from the LRT were reported in Table S6 using the standard DESeq2 notation even if the log_2_-fold changes reported in the LRT test are not directly associated to the test.

Since the LRT determines p-values by comparing dispersion between the full and the reduced model and not by a given pairwise comparison, we did not set a threshold based on fold change. The only criterion that was used to select ASVs significantly influenced by sample types was the p-value adjusted using the Benjamini–Hochberg method (namely an adjusted p-value lower than 0.05). A total of 250 ASVs were selected accounting for 50.92% of the total microbial abundance (mean equals to 56.55% with a standard error of 2.11%). Variance stabilizing transformation (VST, implemented in the vst function of the DESeq2 package) was then used to transform counts into approximately homoskedastic data for clustering. Since abundances of microbial ASVs inferred through amplicon sequencing may have extremely different ranges of values–they reflect actual differences in abundance throughout the community where some species are extremely common and some others can be extremely rare–transformed counts (VST) were also centered and scaled by subtracting the mean of each ASV and dividing results by their standard deviation (a process also known as standardization or z-score transformation). We used a divisive clustering approach to separate ASVs into groups called DIvisive ANAlysis Clustering (diana clustering, implemented in the diana function of the cluster package (*24*)). To group together ASVs with the same pattern across sample types, the Kendall rank correlation coefficient ($\tau$) was transformed into a distance matrix using the formula:

$$d_{ij}=(1-\tau_{ij})^{2}$$

where $d_{ij}$ is the distance between ASV $i$ and ASV $j$ and $\tau_{ij}$ is the value of the Kendall correlation. Doing so, the distance of positively correlated ASVs ranged between 0 and 1 whereas the distance of negatively correlated ASVs ranged between 4 and 1, emphasizing divergences and similarities across ASVs.

The resulting hierarchical clustering was then split into groups by using the divisive coefficient obtained from diana analysis (0.99). Groups containing less than 15 ASVs (roughly 5% of the significant ASVs detected with the LRT) were discarded to reduce spurious clusters (Figure 4 reported in the main text).

Differences in mean across sample types were tested using the non-parametric Wilcoxon signed-rank test. Clusters were tested using all possible combinations of levels (pairwise test) and significance was reported using letters (Figure S7 panel a). P-values were adjusted using the Benjamini–Hochberg method (Figure S7 panel b).

### Enrichment analysis of taxa and functions

To identify enriched/depleted taxa in the clusters produced we used the hypergeometric distribution. Since bacterial lineages were reported at different taxonomic levels (namely, from Domain to Genus/Species), we tested enriched taxa at all levels available to provide a full overview of the clusters defined in the previous chapter. The hypergeometric test is based on the hypergeometric distribution to measure the probability of obtaining a given number of success with a specific number of trials in a defined population (where the number of success and failure are known) (*25*). This definition can be applied to a microbial community with known taxonomic composition. Indeed, we tested the probability of having a given number of ASVs assigned to a specific taxon in a group (cluster) of ASVs in respect with the total community. Following this definition, the probability of having $x$ ASVs defined as a specific taxon of interest can be expressed as:

$$P[X=x]=\frac{\binom{m}{x}\binom{n}{k-x}}{\binom{m+n}{k}}$$

where $m$ is the number of ASVs assigned to the taxon of interest in the whole community, $n$ is the number of taxa outside the group that is being tested, and $k$ is the total number of ASVs in the group. Since we don’t know *a priori* if the taxon is over- or under-represented, we calculated the observed probability of drawing an ASVs belonging to that taxon in the group ($p_{obs}$) and we compare it with the expected probability calculated in the whole community ($p_{exp}$):

$$p_{obs}=\frac{x}{k} ; p_{exp}=\frac{m}{n+k}$$

Then, if $p_{obs}>p_{exp}$, we calculated the probability of having an equal or greater (enriched) number of ASVs of the given taxon. Alternatively, if $p_{obs}<p_{exp}$, we calculated the probability of having an equal or lower (depleted) number of ASVs of the given taxon. This probability represents the p-value of the test obtained using the phyper function of the stats package following the definition reported below:

$$P_{hyper}=\left\{ \begin{matrix} \sum_{x=q}^{k} P[X=x] ; P_{hyper}=P[X\geq x] & \mathrm{if} p_{obs}>p_{exp} \\ \sum_{x=1}^{q} P[X=x] ; P_{hyper}=P[X\leq x] & \mathrm{if} p_{obs}<p_{exp} \end{matrix} \right.$$

We express the magnitude of the enrichment/depletion of a given taxon in a specific group using the log_2_ of fold changes between $p_{obs}$ and $p_{exp}$ (LFC) so that positive values correspond to enriched taxa and negative values correspond to depleted taxa. Obtained p-values were corrected using the Benjamini–Hochberg method to reduce the number of false positives (Table S7). Differently from LRT analysis described above–where we detected the effect of a factor on the overall community without directly comparing the levels of the factor–here we compared the observed probability of having a given number of ASVs assigned to the taxon of interest in a cluster against the same probability calculated in the whole population, hence, we selected significant taxa using both p-value and LFC. Only taxa with a p-value lower than 0.05 and with an LFC value higher than 1 (or less than -1 for depleted taxa) were considered significantly enriched/depleted and used for subsequent analyses.

Similarly to taxonomic enrichment, we detected gene enrichment by using hypergeometric test on gene ontology terms (GO terms) assigned to genes inferred by the PICRUSt2 pipeline (*26*). Enzyme Commission numbers were converted into GO terms following the mapping file available at: <http://www.geneontology.org/external2go/ec2go>. We detected over- and under-represented terms by weighting the number of GO terms found in the inferred genome of ASVs in each group based on genome copy number provided by PICRUSt2 (“genome_function_count” of stratified output). Following the definitions reported above, $m$ represents the number of genes with a given GO term in the whole community, $n$ is the number of genes with the same GO term in the inferred genome content of ASVs outside the group that is being tested, and $k$ is the size of the genome content of ASVs included in the group. Terms significantly enriched or depleted were considered to be shared between ASV groups only if they reported the same effect in the two groups considered (both enriched or both depleted). The number of shared terms between groups was reported in Table S8 whereas the results of gene enrichment analysis were reported in Table S9.

### Tables

**Table S1: Number of samples sequenced for 16S rRNA gene and ITS1 region.** Column one reported the sample id of each condition whereas column two reported the biological matrix sampled (MG, midgut; HG, hindgut; GO, gonads; GI, gills; L, litter; S, soil; W, water; WD, water debris). The sampling site was reported in column three together with the crab’s number (reported in column four). Samples with the same crab’s number within the same sample site are sample coming from the same specimen (experimental unit)–this information was not reported for environmental samples which, in this study, did not share a common experimental unit. The number of sample sequenced was reported in column five and six for 16S rRNA and ITS1 amplicons, respectively. The last column of the table identifies the technical replicates used for ITS1 validation.

**Table S2: Accuracy and correlation between replicates.** Sample IDs were reported in the first column whereas crab’s organs were reported in the second one. The number of fungal ASVs detected in all organs (GI, gills; GO, gonads; HG and MG, hind- and midgut) was reported in the “N” column. Spearman’s rank correlation coefficient was reported in the $\rho$ column whereas accuracy was reported in the last column of the table. Other columns report values used to calculate the accuracy as explained in the “Fungal amplicon sequence variant evaluation” section.

| Sample ID | Organ | N | $\rho$ | TP | TN | FP + FN | Accuracy |
| --- | --- | --- | --- | --- | --- | --- | --- |
| GI_SH_4 | GI | 82 | 0.8 | 10 | 66 | 6 | 0.93 |
| GI_SH_9 | GI | 82 | 0.67 | 5 | 71 | 6 | 0.93 |
| GO_SH_8 | GO | 82 | 1 | 2 | 60 | 20 | 0.76 |
| GO_TKP_7 | GO | 82 | 0.97 | 11 | 70 | 1 | 0.99 |
| HG_SH_1 | HG | 82 | 1 | 5 | 71 | 6 | 0.93 |
| HG_SK_6 | HG | 82 | 0.98 | 12 | 59 | 11 | 0.87 |
| MG_SH_3 | MG | 82 | 0.87 | 6 | 70 | 6 | 0.93 |
| MG_SK_11 | MG | 82 | 1 | 4 | 71 | 7 | 0.91 |

**Table S3: Microbial diversity of samples included in the study.** Microbial diversity was calculated considering counts from both 16S rRNA and ITS1 amplicon sequencing. Sample IDs were reported in the first column (called ID) whereas sample types and sampling sites were reported in the second and the third ones, respectively. The number of ASVs detected in each sample (also known as the “Richness”) was reported together with the number of singletons and doubletons detected (defined as the number of ASVs with a count of one and two across all samples, respectively). The last two columns of the table reported the inverse Simpson index and the Good’s coverage for all samples (for additional information on how these indices were computed see “Microbial diversity within and between sample types” section).

**Table S4: Variance explained by crab, sample type, and sampling site computer with permutational multivariate analysis of variance (PERMANOVA).** The variance explained by each factor (also called R squared) was reported separately for 16S and ITS1 amplicons. The type columns reports if the PERMANOVA was computed on crab’s samples (midgut, hindgut, gills, and gonads) or on environmental samples (litter, soil, water, and water debris). The crab was included in the model to account for variability across different animals collected. Asterisks were used to report significance based on p-values (*, p-value < 0.05; ** p-value < 0.01). The proportion of variance unexplained by the factor considered was reported in the Residual column.

| Amplicon | Type | Crab | Sample Type | Site | Residual |
| --- | --- | --- | --- | --- | --- |
| 16S | Crab’s organs | 0.203** | 0.260** | 0.043** | 0.494 |
|  | Environmental samples | - | 0.160** | 0.088** | 0.752 |
| ITS | Crab’s organs | 0.226 | 0.053** | 0.055** | 0.666 |
|  | Environmental samples | - | 0.174** | 0.132** | 0.694 |

**Table S5: Analysis of variance (ANOVA) on euclidean distances around centroids.** The homogeneity of dispersion across groups was tested using ANOVA on euclidean distances from group centroids. P-values were reported in the table using asterisks to report statistical significance (*, p-value < 0.05; ** p-value < 0.01).

| Amplicon | Type | Sample Type | Site |
| --- | --- | --- | --- |
| 16S | Crab’s organs | 0.672 | 0.012* |
|  | Environmental samples | 0.206 | 0.979 |
| ITS | Crab’s organs | 0.067 | 0.024* |
|  | Environmental samples | 0.087 | 0.479 |

**Table S6: Results of the likelihood ratio test (LRT) on sequence variants in different sample types.** All columns returned from the LRT were reported even if the “log2FoldChange” column is not representative of a given contrast in this type of test. Columns are: asv, the name of the ASV tested; baseMean, the mean overall abundance of the each ASV tested; log2FoldChange, the logarithm of the ratio between two levels of the factor considered (since in LRT no contrast is provided, this column is not directly associated with the actual hypothesis test); lfcSE, the standard error of the log2FoldChange; stat, the difference in deviance between the reduced model and the full model; pvalue, the pvalue obtained by comparing stat to a chi-squared distribution; padj, adjusted p-values using the Benjamini–Hochberg method.

**Table S7: Taxa enriched/depleted in the clusters defined base on LRT test.** Columns report the number of ASVs found in the group of interest and in the whole community as used in the hypergeometric test. In details: Amplicon, the amplified amplicon used for barcoding (16S stands form 16S rRNA gene V3-V4 regions and ITS stands for the ITS1 region); Level, taxonomic level considered; Taxon, the name of the taxon tested; Cluster, the cluster in which the enrichment/depletion was tested; x, the number of ASVs assigned to the taxon considered in the cluster of interest; k, the number of ASVs in the cluster considered; n, the number of ASVs outside the cluster; m, the number of ASVs assigned to the given taxon in the whole community; p_obs, the probability of having an ASVs assigned to the taxon of interest in the cluster; p_exp, the probability of having an ASVs assigned to the taxon of interest in the community; log2.fold.enrichment, $log_{2}$ of the ratio between p_obs and p_exp; p.value, the p value; adj.p.value, p value adjusted following the Benjamini–Hochberg method.

**Table S8: Number of shared GO terms between clusters.** Columns C1, C2, and C3 represent the ASV groups (C1, cluster 1; C2, cluster 2; C3, cluster 3), if an “X” is present then GO terms are included in the group. The number of GO terms enriched/depleted in each group is reported in the last two columns. The total number of terms detected in the inferred genomes of ASVs assigned to each cluster was marked with “Tot” and reported in the last three rows of the table.

| C1 | C2 | C3 | Depleted | Enriched |
| --- | --- | --- | --- | --- |
| X |  |  | 18 | 200 |
|  | X |  | 12 | 76 |
|  |  | X | 19 | 204 |
| X | X |  | 1 | 51 |
| X |  | X | 2 | 71 |
|  | X | X | 1 | 67 |
| X | X | X | 0 | 37 |
| Tot |  |  | 21 | 359 |
|  | Tot |  | 14 | 231 |
|  |  | Tot | 22 | 379 |

**Table S9: Gene ontology terms enriched/depleted in ASV clusters.** Columns report the number of genes found in the inferred genome of ASVs included in the group of interest as well as in the whole community as used in the hypergeometric test. Values were weighted using the inferred copy number as obtained by PICRUSt2 (“genome_function_count”). Columns correspond to: GO, gene ontology term; Cluster, the group of ASVs considered; x, the number of genes classified with the GO term reported in the cluster of interest; k, the number of GO terms in the cluster considered; n, the number of GO terms outside the cluster; m, the number of GO term detected in the inferred genome of the ASVs in the whole community; p_obs, the probability of having a gene assigned to the GO term of interest in the ASV cluster; p_exp, the probability of having a gene assigned to the GO term of interest in the whole community; log2.fold.enrichment, log_2_ of the ratio between p_obs and p_exp; p.value, the p value; adj.p.value, p value adjusted following the Benjamini–Hochberg method.

### Figures

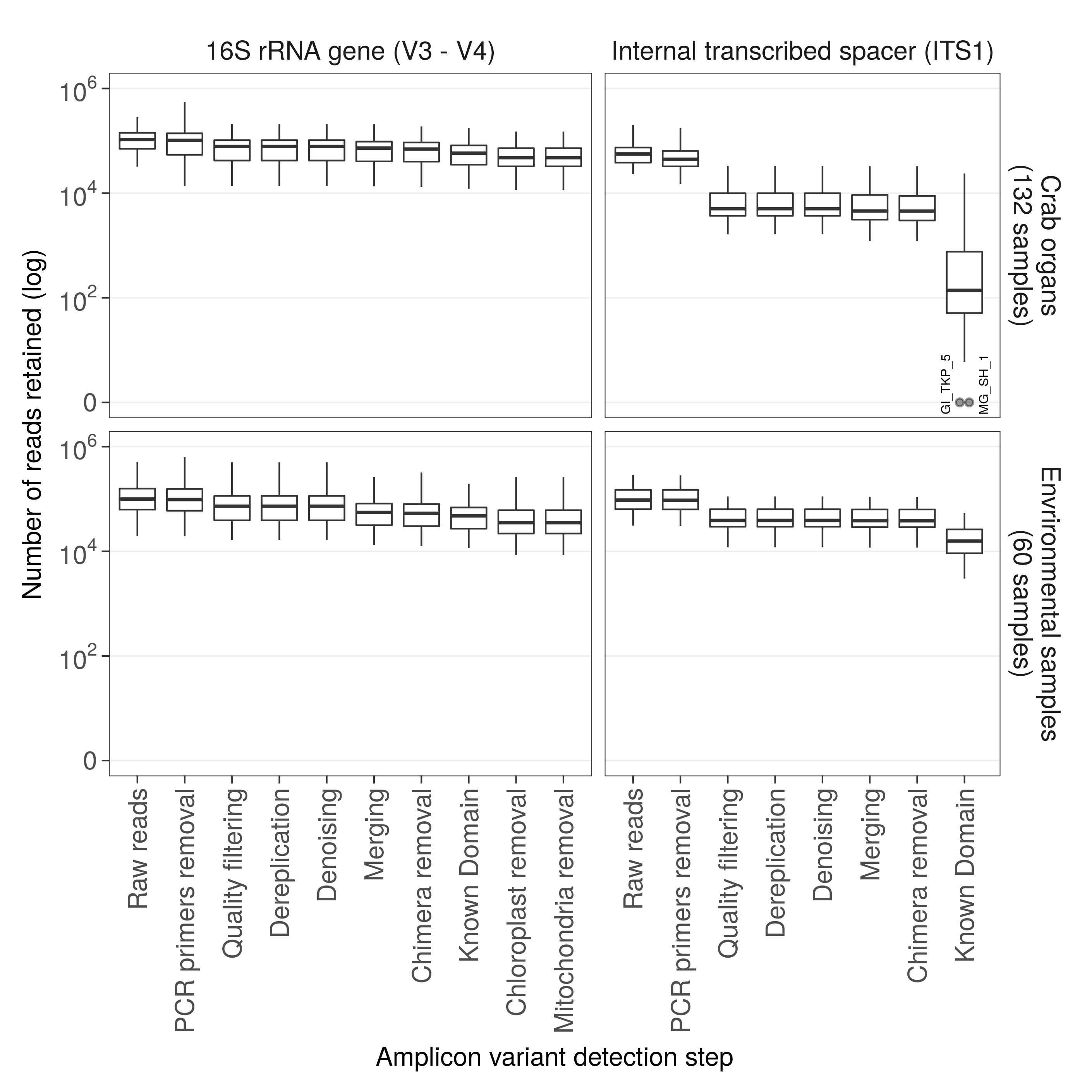

**Figure S1: Number of reads retained after each step of the DADA2 workflow.** The number of reads processed and retained after each step of the DADA2 workflow (x axis) is reported in the y axis. Results obtained from 16S rRNA gene the ITS1 region amplicons were reported in two different panels and divided according to the sample type (crab’s organs or environmental samples). The sample ID of two fungal samples (ITS1 region) which resulted in zero counts after DADA2 pipeline were reported in the plot.

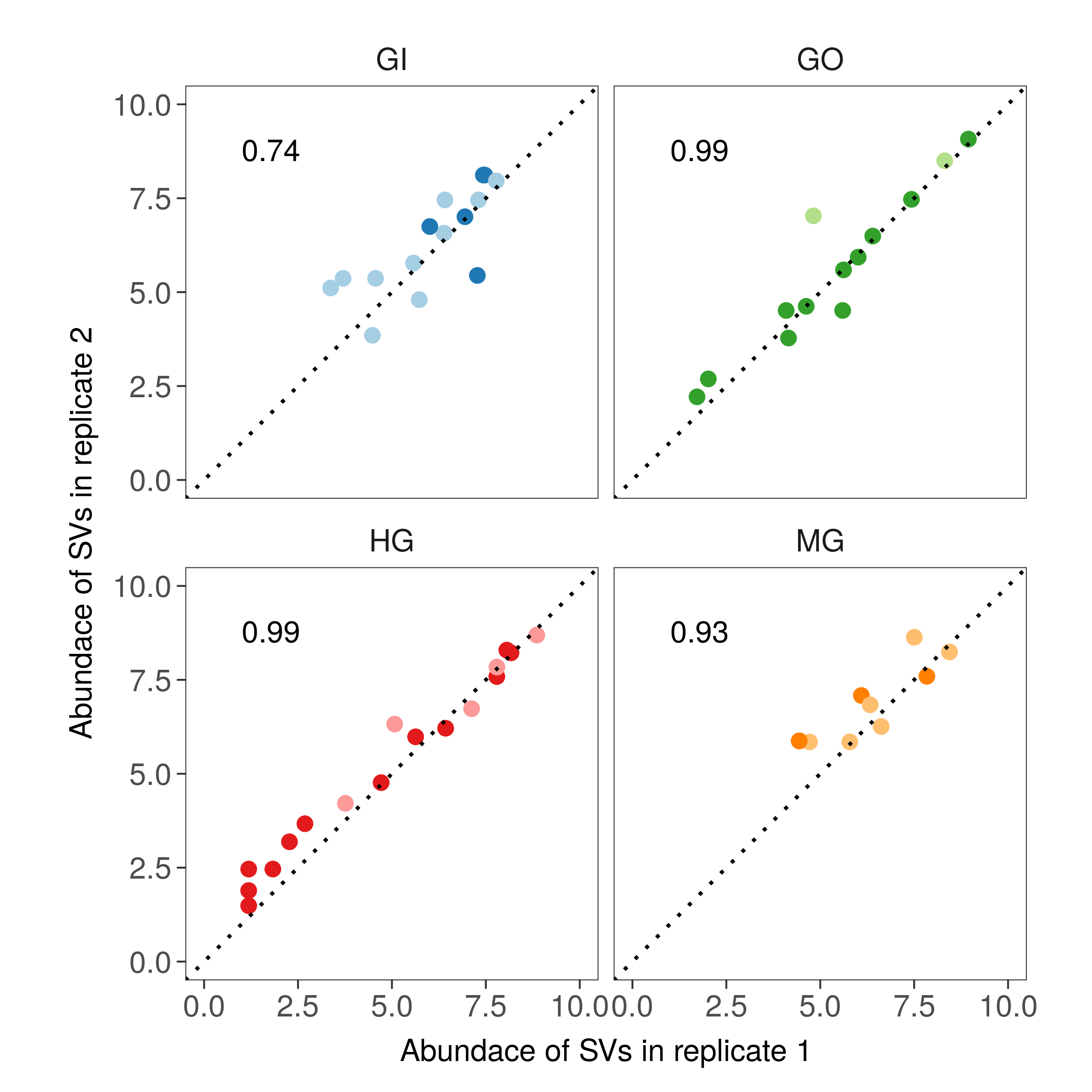

**Figure S2: Counts of SVs for each replicate in different crab’s organs.** The abundance of every ASV is reported for the eight pairwise combinations using the log2-scale (2 contrast and four crab’s organs). Colors represent crab’s organs (GI, gills, blue; GO, gonads, green; HG, hindgut, red; MG, midgut, orange) whereas samples were reported using different shades of colors (two samples per organ, one darker than the other). Dotted lines represent a perfect correlation (namely a line with slope equal to one and intercept equal to zero) whereas the average value of the Spearman’s correlation coefficient was reported at the top left of each panel.

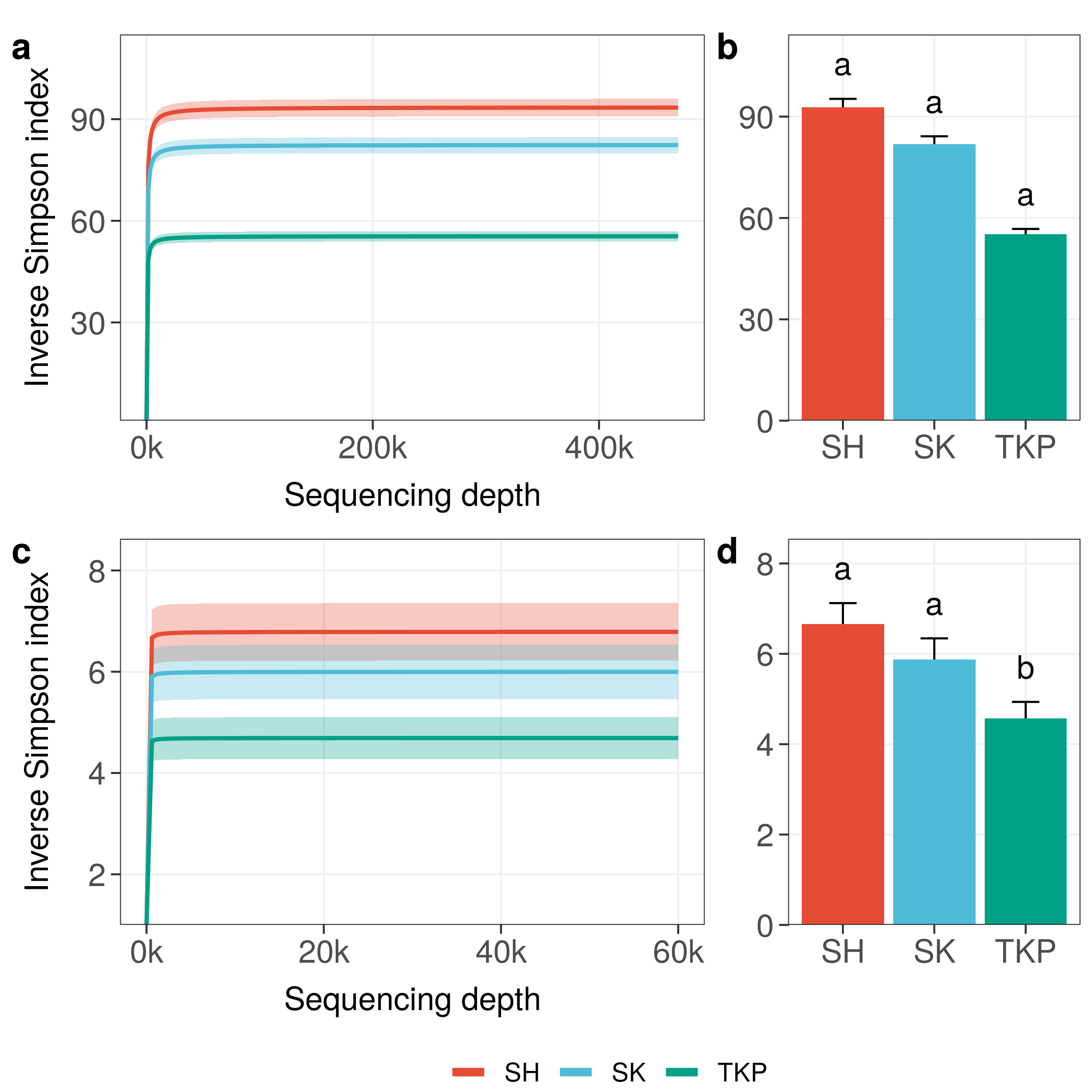

**Figure S3: Microbial diversity in different sampling sites.** The average of the inverse Simpson index was reported with increasing sampling effort for all sampling sites (SH, Shui Hau; SK, Sai Keng; TKP, To Kwa Peng). Interpolated and extrapolated diversity was reported in panel a and c (16S rRNA gene and ITS1 region, respectively), whereas observed diversity was reported in panel b and d (16S rRNA gene and ITS1 region respectively). Significant differences in microbial diversity (Wilcoxon non-parametric test) were reported using lowercase letters (panel b and d) whereas colors and acronyms on the x-axis correspond to different sample sites. If two means were significantly different, all letters on top of the two boxes must be different; if two means were equal, at least one letter must be the same.

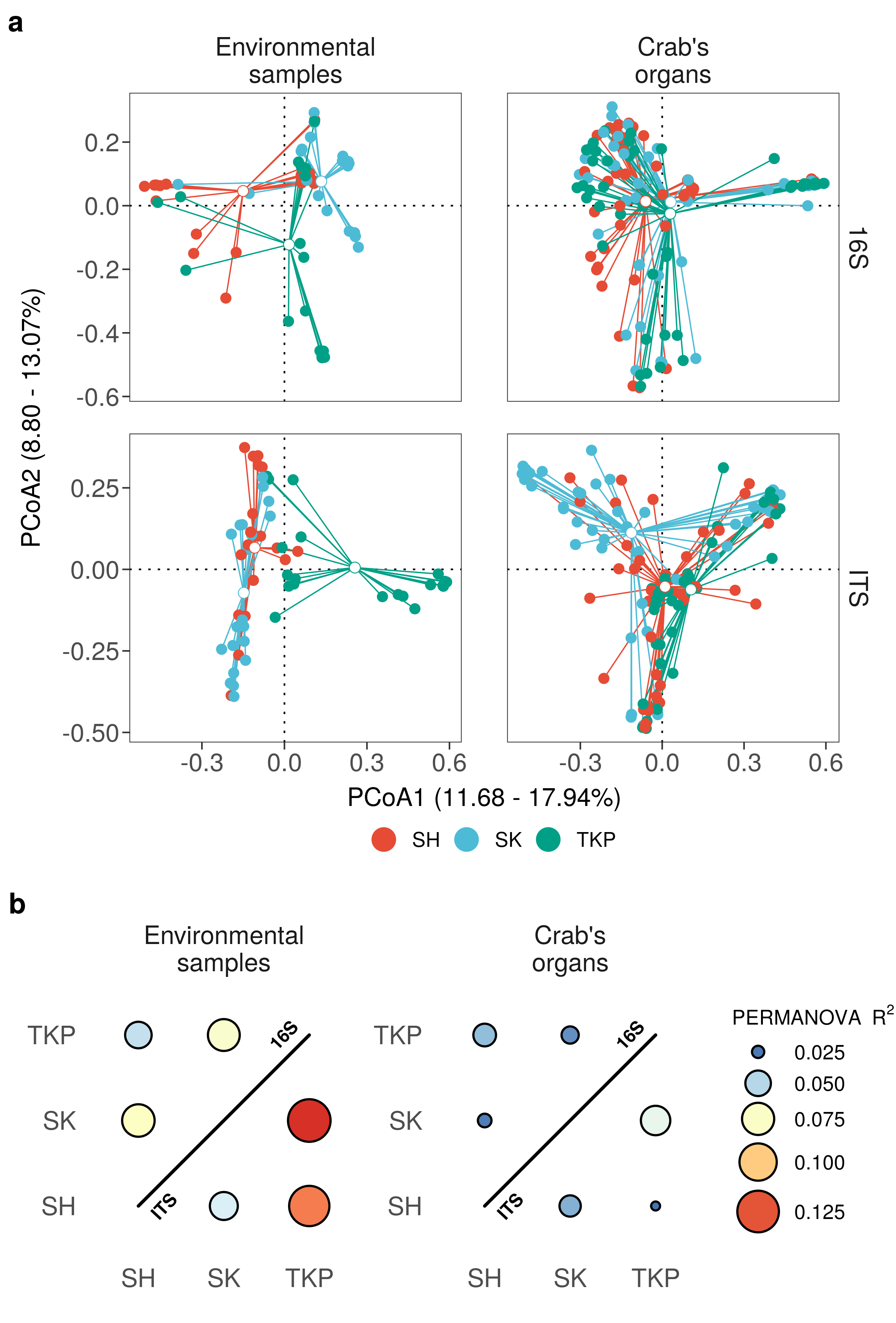

**Figure S4: Microbial distribution according to Bray-Curtis distance.** a) Principal coordinates analyses on Bray-Curtis distances inferred form 16S rRNA and ITS1 metabarcoding. Environmental samples and crab’s organs (top side of the panels) were reported separately as distribution obtained with the two markers reported (right side of each panel). Solid-colored points represent different samples whereas white-filled points represent centroids. The variance of the objects along each axis has been reported between squared brackets (from the lowest to the highest) and different sample types were reported using different colors (TKP, To Kwa Peng; SK, Sai Keng; SH, Shui Hau). b) Permutational analysis of variance on ordinations reported in panel a. For each pair of organs and environmental samples a permutational analysis of variance was performed. R-squared values of significant contrasts were reported for both 16S (upper triangle) and ITS (lower triangle) counts.

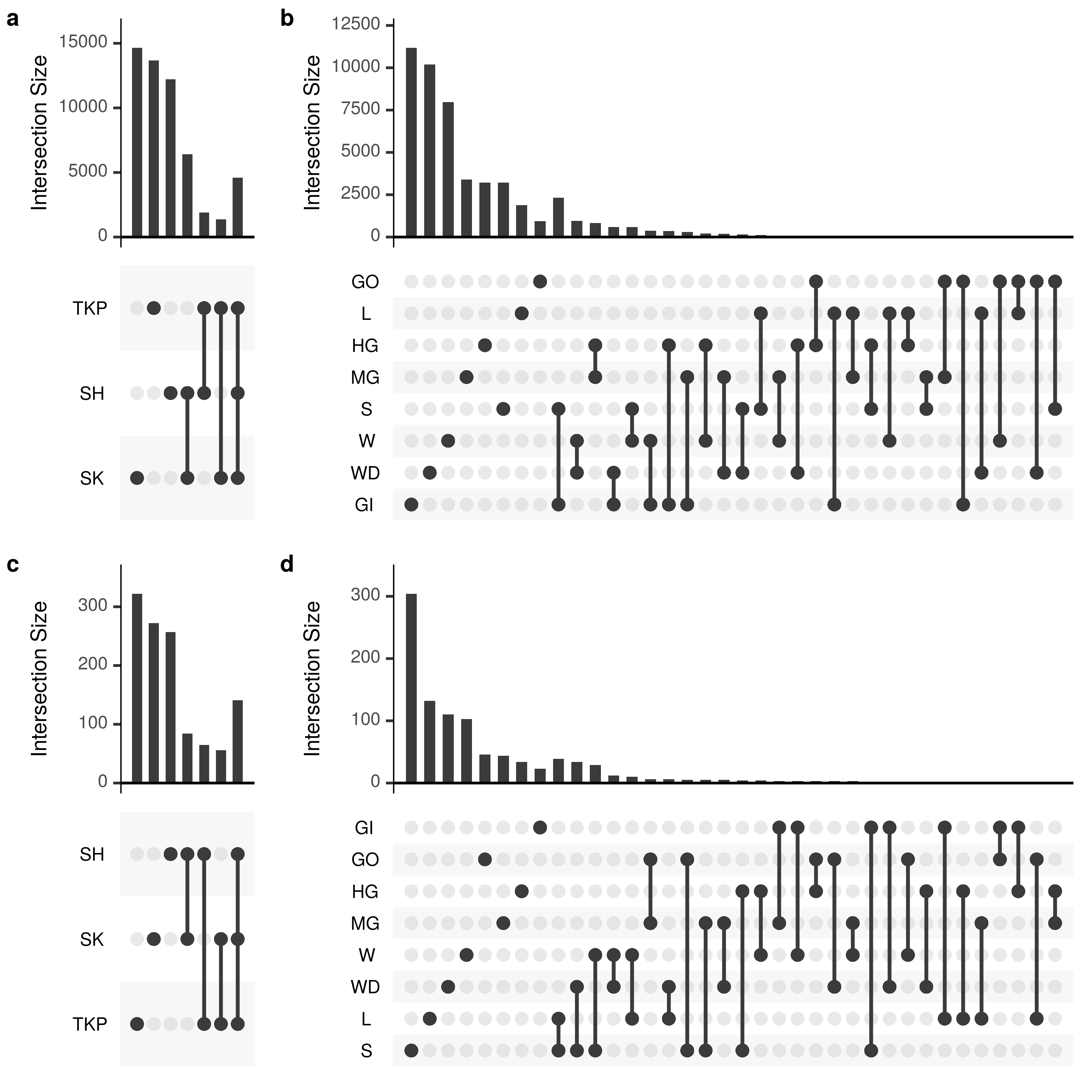

**Figure S5: Number of ASVs shared across sampling sites and sample types.** The number of ASVs in common between different set intersections was reported following the upset representation. Panels a and b report the number of shared ASVs obtained from 16s rRNA amplicon sequencing whereas panels b and d report those obtained form ITS1 sequencing. Set intersections were displayed in a matrix layout where each row is a different site (panels a and c) or sample type (panels b and d) and each column corresponds to a different intersection. The number of ASVs contained in each intersection is reported using bars on top of the intersection considered. Intersections are mutually exclusive so that if an ASV is present in a given intersection it is excluded from the others. Sites and sample types were abbreviated as follows: TKP, To Kwa Peng; SK, Sai Keng; SH, Shui Hau; MG, midgut; HG, hindgut; GO, gonads; GI, gills; L, litter; S, soil; W, water; WD, water debris.

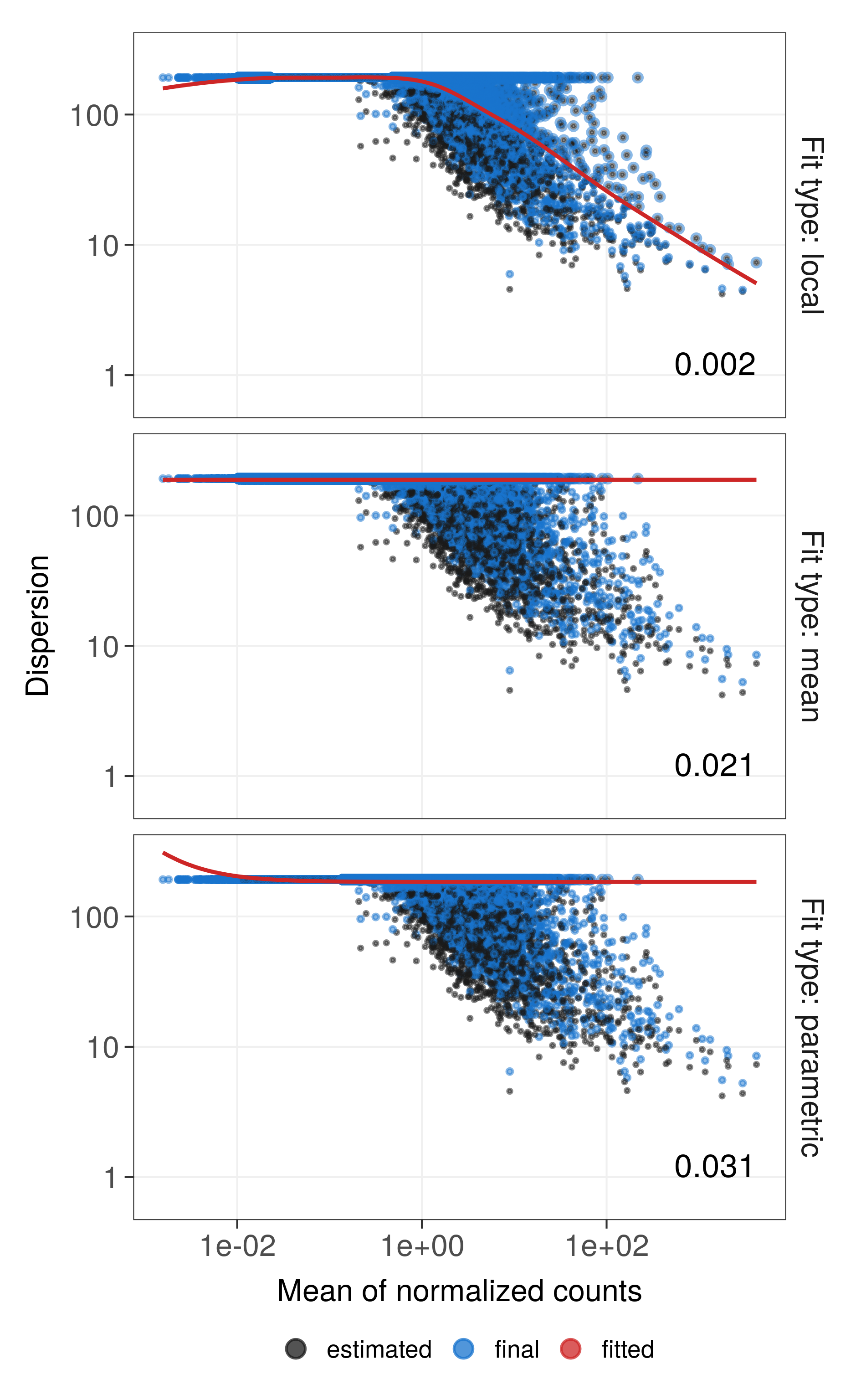

**Figure S6: DESeq2 model selection based on the median of residuals on the log scale.** Per-ASV dispersion estimates (x-axis) were plotted together with the mean of normalized counts obtained from DESeq2 (y-axis). Black dots represent the maximum-likelihood estimates (MLEs) for each ASVs using only the respective data, fitted models were reported using a red line, and the final estimates used fo testing were reported using blue dots. Outliers were highlighted with blue circles. All fit types available were tested and the median absolute log residuals obtained were reported in the bottom right corner of each plot.

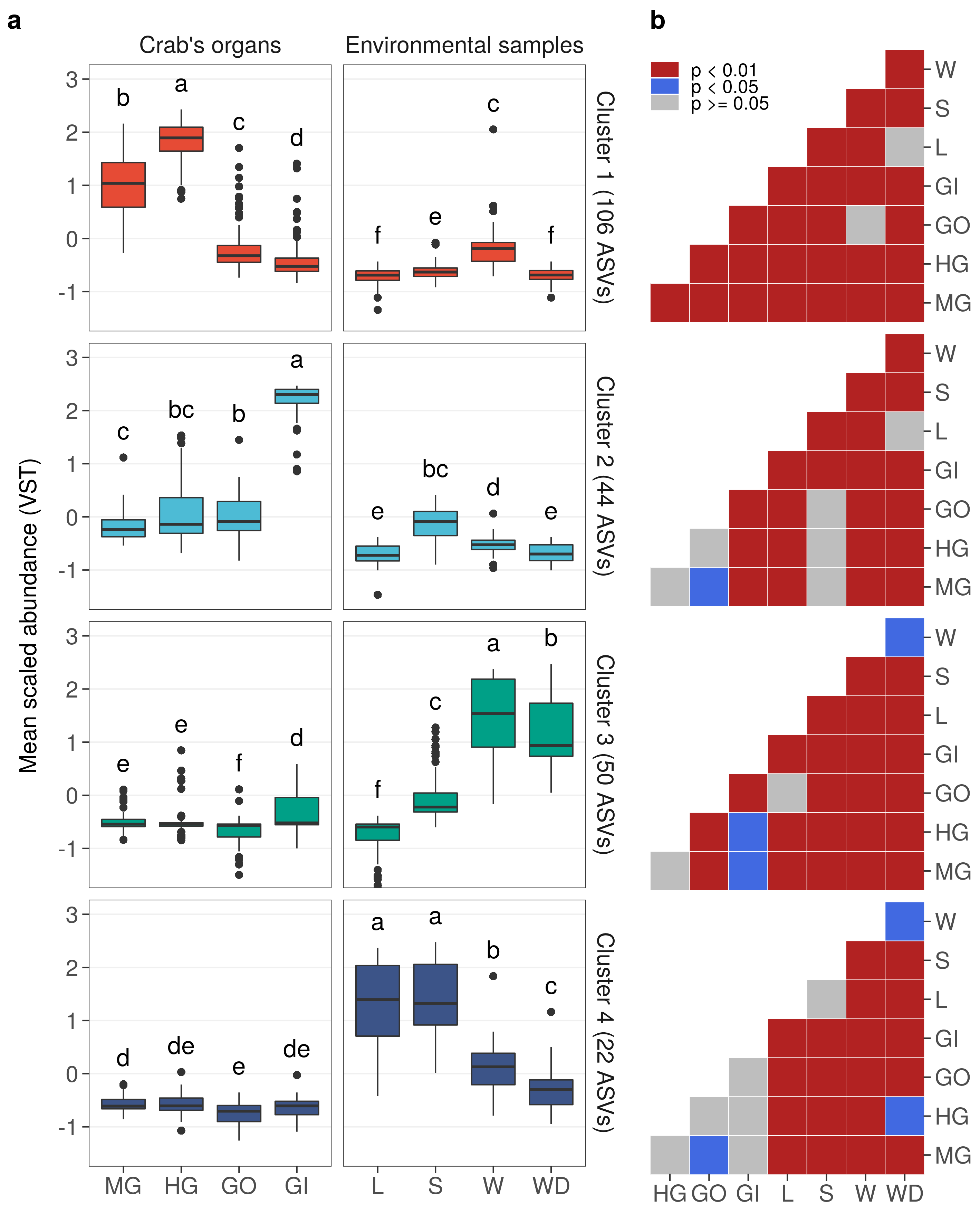

**Figure S7: Differences across sample types in the four clusters detected with divisive hierarchical clustering.** a) The abundance values of all ASVs included in clusters was reported using the box and whisker visualization where: the interquantile range (IQR, the space between Q_3_ and Q_1_) is represented by the “box”, the “whiskers” extend to the most extreme data point which is no more than 1.5 times the IQR away from the box, and the median (Q_2_) is reported with a black horizontal line. Outliers, namely those observations exceeding 1.5 times the IQR, were reported using black dots. Lowercase letters on top of each box represent significant differences across sample types: if two means were significantly different, all letters on top of the two boxes must be different; if two means were equal, at least one letter must be the same. Crab’s organs and environmental samples were reported in two different panels (vertically) but all pairwise contrasts were included in the analysis. b) Adjusted p-values between pairwise contrasts. All contrasts were tested using Wilcoxon signed-rank test and results were reported as an heatmap where red and blue cells represent two levels of significance (p < 0.01 and p < 0.05, respectively) whereas gray cells indicate non statistical significance (p > 0.05).
